# Autoimmunity Risk Gene IRGM is a Master Negative Regulator of Interferon Response by Controlling the Activation of cGAS-STING and RIG-I-MAVS Signaling Pathways

**DOI:** 10.1101/815506

**Authors:** Kautilya Kumar Jena, Subhash Mehto, Parej Nath, Nishant Ranjan Chauhan, Rinku Sahu, Tapas Kumar Nayak, Saroj Kumar Das, Kollori Dhar, Pradyumna Kumar Sahoo, Krushna C Murmu, Saikat De, Ankita Datey, Punit Prasad, Soma Chattopadhyay, Swati Chauhan, Santosh Chauhan

## Abstract

Activation of type 1 interferon response is extensively connected with the antiviral immunity and pathogenesis of autoimmune diseases. Here, we found that IRGM, whose deficiency is linked with the genesis of several autoimmune disorders, is a master negative regulator of the interferon response. Mechanistically, we show that IRGM interacts with nucleic acid sensor proteins, including cGAS and RIG-I, and mediates their autophagic degradation to restrain activation of interferon signaling. Further, IRGM maintains mitophagy flux, and its deficiency results in the accumulation of defunct leaky mitochondria that releases cytosolic DAMPs triggering activation of interferon responses via cGAS-STING and RIG-I-MAVS signaling axis. Due to an enduring type 1 IFN response in IRGM-deficient cells and mice, they were intrinsically resistant to infection of the Japanese Encephalitis virus, Herpes Simplex virus, and Chikungunya virus. Altogether, this study defines the molecular mechanisms by which IRGM maintains interferon homeostasis and protects from autoimmune diseases. Further, it identifies IRGM as a broad therapeutic target for defense against viruses.

**Graphical Abstract:** 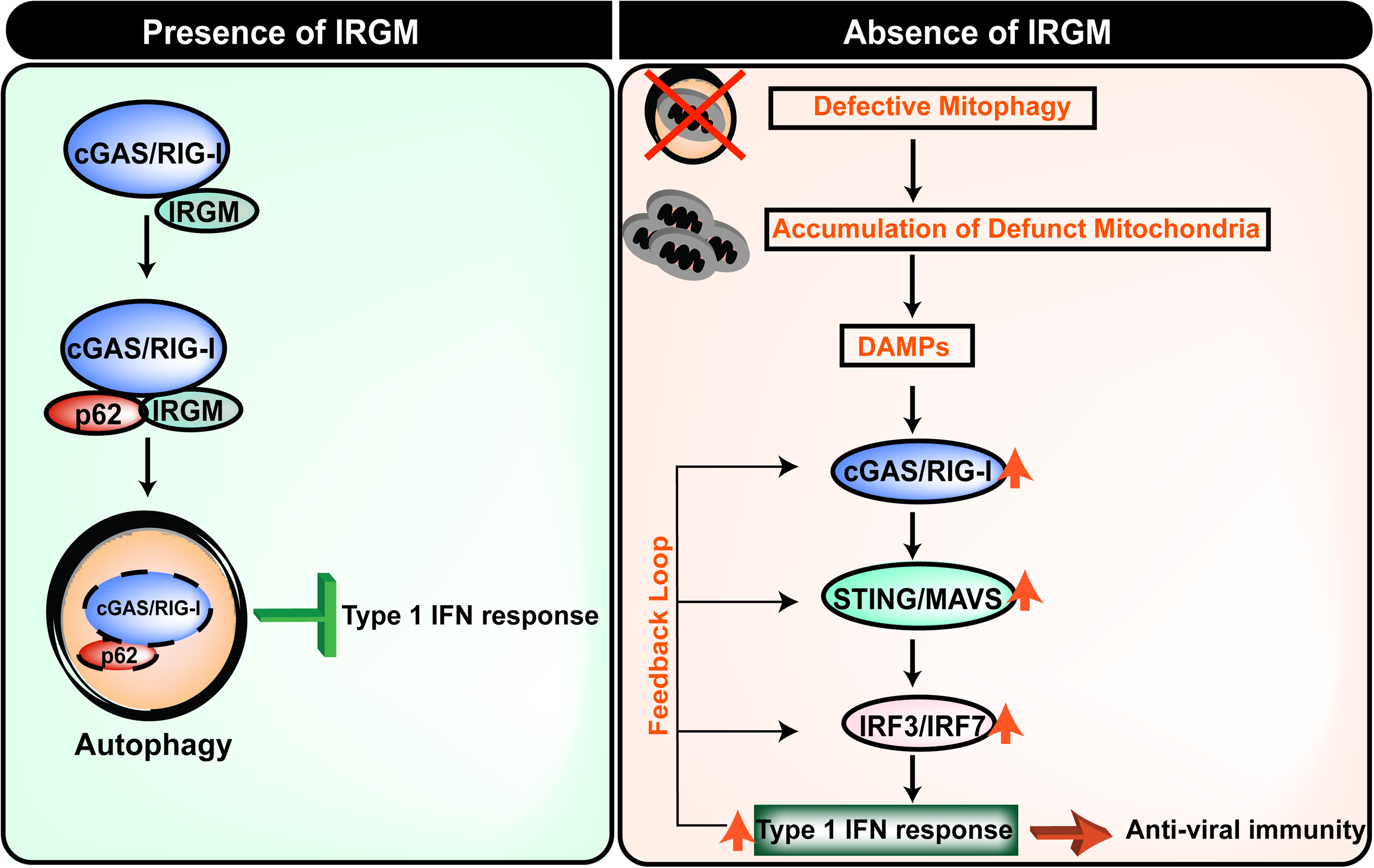

## Introduction

Our understanding of the activation of innate immune systems upon exposure to hostile conditions such as microbial infection has grown exponentially (Akira et al., 2006; Goubau et al., 2013; Takeuchi and Akira, 2010). However, how the innate immune pathways are controlled under steady-state conditions are not well defined. In particular, the mechanisms by which negative regulators of innate immune system restrain the aberrant immune activations under basal conditions need to be illustrated to understand the genesis of spontaneous inflammatory diseases, including autoimmune disorders. The type I interferon (IFN) response, where one side is first-line of defense against invading pathogens (esp. viruses), on the other side, its uncontrolled activation can lead to several autoimmune diseases (Crow et al., 2019; Di Domizio and Cao, 2013; Psarras et al., 2017). A fine homeostatic balance of type I interferons needs to be maintained to avoid autoimmune diseases, including interferonopathies (Crow et al., 2019; Di Domizio and Cao, 2013; Lee-Kirsch et al., 2016; Niewold, 2014). The knowledge of the master switches and the mechanisms that suppress the type I IFN response will be beneficial for generating therapeutics against autoimmune diseases.

The pattern recognition receptors (PRR’s) senses external (pathogen and PAMPs, pathogen-associated molecular patterns) and internal (DAMPs, danger-associated molecular patterns) cellular threats and mount a strong innate immune response that includes the production of pro-inflammatory cytokines (Roers et al., 2016; Takeuchi and Akira, 2010). The presence of PAMPs or DAMPs in the cytosol are sensed by cytosolic PRRs such as RIG-I like receptors (RLR) and NOD-like receptors (NLRs) and also by DNA and RNA sensors such cGAS, IFI16 and ZBP-1 (Kuriakose and Kanneganti, 2018; Radoshevich and Dussurget, 2016; Roers et al., 2016; Unterholzner et al., 2010; Wu and Chen, 2014). RIG-I or MDA5 senses dsRNA species and activates adaptor protein MAVS, which then acts as a platform for activation of TBK1 and IRF3/IRF7 transcription factors (Hornung et al., 2006; Kato et al., 2006; Reikine et al., 2014). These transcription factors then translocate to the nucleus to increase the production of type I interferons. Similarly, DNA sensor cGAS upon sensing dsDNA of viral, mitochondrial or genomic origin activates adaptor protein STING leading to activation of TBK1-IRF3/IRF7 axis for type I interferon production (Li et al., 2013; Mackenzie et al., 2017; Roers et al., 2016; Rongvaux et al., 2014; Sun et al., 2013; West et al., 2015). The interferons thus produced can activate the JAK-STAT1/2 signaling pathway leading to the production of interferon-stimulated genes (ISGs), which are the powerful effector proteins with a varied function in innate immunity, including anti-viral/bacterial response (Ivashkiv and Donlin, 2014; Roers et al., 2016). The imbalance in all of these signaling pathways has been strongly linked with autoimmunity (Di Domizio and Cao, 2013; Gray et al., 2015; Kato and Fujita, 2015; Louis et al., 2018; Rice et al., 2014)

IRGM (Irgm1) deficiency is genetically and functionally associated with several inflammatory and autoimmune diseases including ankylosing spondylitis, autoimmune thyroid diseases, Graves’ disease, Sjogren’s syndrome, Crohn’s disease, experimental autoimmune encephalomyelitis, Hepatic steatosis, NAFLD (non-alcoholic fatty liver disease) and severe sepsis (Azzam et al., 2017; Bellini et al., 2017; Kimura et al., 2014; Lin et al., 2016; Parkes et al., 2007; Xia et al., 2017; Xu et al., 2010; Yao et al., 2018). Recently, in knock out mouse model, Irgm1 (mice orthologue of IRGM) was shown to control autoimmunity (Azzam et al., 2017). They show that naive Irgm1 knock out mice in the germ-free conditions displayed the hallmarks of Sjogren’s syndrome, an autoimmune disorder characterized by lymphocytic infiltration of exocrine tissues (Azzam et al., 2017). The presence of IRGM/Irgm1 in humans and mice is shown to be largely protective against autoimmune disorders. The connections between IRGM and systemic autoimmune diseases argues a gigantic role of IRGM in innate immune homeostasis. The molecular mechanism by which human IRGM controls innate immune homeostasis in steady-state conditions remains completely undetermined.

A large number of accumulating evidence suggests that autophagy-mediated clearance of defunct mitochondria is a powerful mechanism to keep the inflammation under check (Oka et al., 2012; Sliter et al., 2018; Tal et al., 2009; Xu et al., 2019). Autophagy deficiency results in the accumulation of dysfunctional mitochondria that are the major source of DAMPs for activation of cGAS-STING and RIG-I/MAVS signaling pathways. Activations of these pathways lead to robust induction of interferon response resulting in antiviral response or autoimmune diseases (Gkirtzimanaki et al., 2018; Sliter et al., 2018; Tal et al., 2009; Xu et al., 2019). Others and we have found that IRGM is a key autophagy protein that plays a significant role in anti-bacterial autophagy and autophagy of inflammasomes (Chauhan et al., 2015; Kumar et al., 2018; Mehto et al., 2019; Singh et al., 2006; Singh et al., 2010). IRGM was also shown to present over mitochondria, and overexpression of IRGM induces mitochondrial fission, followed by its depolarization (Singh et al., 2010). However, it is completely undetermined whether IRGM deficiency perturbs mitophagy and affects the downstream innate immune signaling pathways.

This study uncovers that under homeostatic conditions, IRGM is a master suppressor of type I IFN response. The mRNA sequencing whole transcriptome analysis in human cells and mice shows that IRGM controls the expression of almost all major ISG’s. Mechanistically, we show that IRGM suppresses IFN signaling by mediating p62-dependent autophagic degradation of cGAS, RIG-I, and TLR3. Further, we found that IRGM is critical for the removal of damaged mitochondria by autophagy. Thus, IRGM deficiency results in defective mitophagy, accumulation of dysfunctional mitochondria, and enhanced mitochondrial DAMPs that stimulate cGAS-STING and RIG-I-MAVS axis to drive robust activation of type I IFN response. In a nutshell, IRGM maintains the homeostatic balance of IFNs and ISGs by (**1**) mediating degradation of DNA/RNA sensor proteins and (**2**) by maintaining efficient mitophagy and healthy mitochondria. Furthermore, we show that due to heightened type 1 IFN response in IRGM depleted cells and mice, they are intrinsically resistant to infection of both DNA and RNA viruses.

## Results

### IRGM is a master suppressor of the interferon response

To understand the role of IRGM in innate immune homeostasis and autoimmunity, we performed RNA sequencing (RNA-seq) experiments with (**1**) control and IRGM shRNA knockdown (hereafter, **IRGM KD, Supplementary Figure 1A**) human HT29 colon epithelial cell line, (**2**) wild type (***Irgm1^+/+^***) and *Irgm1* knockout mice (***Irgm1^-/-^***) bone marrow-derived macrophages (BMDM’s), and (**3**) *Irgm1^+/+^* and *Irgm1^-/-^*brain tissues.

The gene ontology (GO) based pathway analysis was performed using Ingenuity pathway analysis (IPA, https://analysis.ingenuity.com/), Reactome pathway analysis (Fabregat et al., 2018), and Metascape pathway analysis (Tripathi et al., 2015) with genes upregulated (1.5 folds, p<0.05, n=3) in IRGM KD HT29 cells. In all of these analyses, the top-enriched pathways were the induction of innate/adaptive immune systems and inflammatory signaling/responses (**Figure 1A, Supplementary Figure 1B and 1C**) indicating that the primary function of human IRGM under steady-state conditions is to control the cellular inflammation and immunity. A closer look at the genes and the pathways that are upregulated suggest that IRGM deficiency results in the induction of interferon responses or the processes/pathways controlled by the interferon responses (**Figure 1A, Supplementary Figure 1B and 1C, Table S1)**. To our surprise, almost all well-known interferon-stimulated genes (ISG’s) including IFI ‘s (interferon-inducible genes), OAS genes (Oligoadenylate synthase’s), ISG15/20, GBP’s (Guanylate-binding protein’s), APOBEC genes (Apolipoprotein B mRNA Editing Catalytic Polypeptide-like), MX genes (myxovirus resistance), MHC class 1 antigen processing and presentation genes, and TRIM’s (tripartite motifs) genes were upregulated upon knocking down IRGM (**Figure 1A, Supplementary Figure 1B and 1C, Table S1**). The Interferome database analysis (Rusinova et al., 2013) using highly stringent parameters shows that ∼45% of the genes (392 out of a total of 890) induced in IRGM KD cells are interferon-regulated (**Figure 1B**). The interferons are the major defense system against viruses, and that is the reason why the “defense response to viruses/microbes” are other top-induced functions in the IPA (**Figure 1C, Table S2)** and Metascape pathway analysis **(Supplementary Figure 1C)**. In IPA, cancer, autoimmunity (Psoriasis, Sjogren’s syndrome, age-related macular degeneration), and other inflammatory disorders were the top diseases associated with IRGM deficiency (**Supplementary Figure 1D, Table S2**). The qRT-PCR was performed with key interferon-inducible genes (*RIG-I, IFI16, MDA5, STAT2, OAS1, MX2, ISG15, TRIM22, and APOBEC3G*) to validate the RNA-seq data (**Figure 1D**). We observed 2 to 1200 folds induction of ISG’s in IRGM deficient cells suggesting that IRGM is a potent inhibitor of interferon response (**Figure 1D**). The RNA-seq data was also validated in human THP-1 monocytic cell line (data present throughout the manuscript), and also the expression of few of the ISG’s was validated in human peripheral blood mononuclear cells (PBMC’s) from three independent human donors, where IRGM was knockdown using siRNA (**Supplementary Figure 1E**).

**Figure 1.**
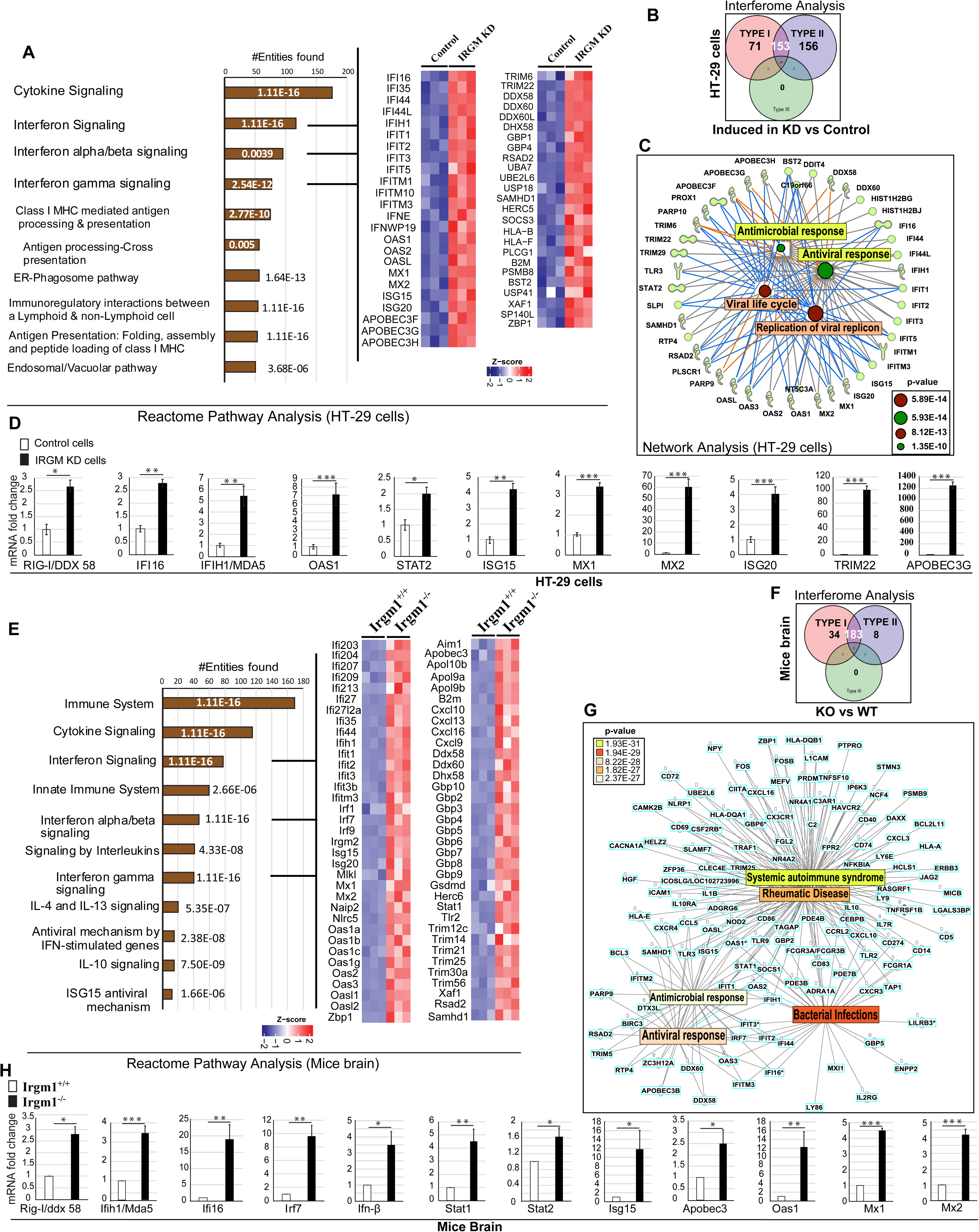
IRGM is a master negative regulator of the interferon response. **(A)** The bar graph represents top 10 biological pathways upregulated in gene ontology (GO) based Reactome pathway analysis using a set of genes induced (1.5 fold, p<0.05, 3 biological replicates) in RNA-seq analysis in IRGM shRNA knockdown HT29 cells compared to control shRNA cells. Heatmaps were generated for sentinel interferon-regulated genes (three biological replicates) using ‘ComplexHeatmap’ library using ‘R’ Bioconductor package where the gene expression matrix was transformed into z-score **(B)** Interferome database analysis with a set of genes induced (1.5 fold, p<0.05, 3 biological replicates) in IRGM shRNA knockdown HT29 cells compared to control shRNA cells. The Venn diagram depicts the total number of upregulated type I and type II IFN-regulated genes in IRGM KD cells. **(C)** Network pathway analysis using IPA. The molecular network of genes connected with top 4 functions associated genes (1.5 fold, p<0.05, 3 biological replicates) upregulated in IRGM knockdown HT29 cells. The complete list is documented in supplementary table S2. **(D)** qRT-PCR validation of RNA-seq data in control and IRGM KD HT29 cells. Mean ± SD, n = 3, *P < 0.05, **P < 0.005, ***P < 0.0005, Student’s unpaired t-test. **(E)** The bar graph represents top 10 pathways upregulated in GO-based Reactome pathway analysis using set of genes induced (1.5 fold, p<0.05, 3 mice each group) in the brain of *Irgm1^-/-^* mice compared to *Irgm1^+/+^*wild type mice. Heatmaps were generated for sentinel interferon-regulated genes (three biological replicates) using ‘ComplexHeatmap’ library using ‘R’ Bioconductor package where the gene expression matrix was transformed into z-score. **(F)** Interferome database analysis with a set of genes induced (1.5 fold, p<0.05) in the brain of *Irgm1^-/-^*mice compared to *Irgm1^+/+^* wild type mice. The Venn diagram depicts the total number of upregulated type I and type 2 IFN-regulated genes in *Irgm1^-/-^* mice brain. **(G)** Network pathway analysis using IPA. The molecular network of genes connected with top 5 functions/diseases associated with genes (1.5 fold, p<0.05) upregulated in *Irgm1* knockout mice brain. The complete list is documented in supplementary table S2. **(H)** The qRT-PCR validation of RNA-seq data in *Irgm1^+/+^*and *Irgm1^-/-^* mice brain. Mean ± SD, n = 3, *P < 0.05, **P < 0.005, ***P < 0.0005, Student’s unpaired t-test.

Next, we performed pathway analysis with RNA-seq data from brain and BMDM’s of *Irgm1^+/+^* and *Irgm1^-/-^* mice (n=3). The reason for performing RNA-seq with brain tissues is that it is a relatively immune-privileged organ and is mostly insusceptible to perturbation in the peripheral immune system due to extraneous irritants and, thus, immune responses are typically cell-intrinsic. There was a remarkable similarity in the upregulated genes and pathways in IRGM-depleted human HT29 cells, the *Irgm1^-/-^* mice brain, and the *Irgm1^-/-^* BMDM’s (**Figure 1A and 1E, Supplementary Figure 1F, 1G and 2A, Table S1)**. Both in the brain and BMDM’s the pathways that were enriched as a response of *Irgm1* knockout were related to cytokine response, interferon signaling/response, and antiviral/microbial response (**Figure 1E, Supplementary Figure 1F, 1G and 2A, 2B)**. Remarkably, >80% of the genes (225 out of 288) that were upregulated in *Irgm1^-/-^* brain and >50% of the genes (314 out of 595) that were induced in *Irgm1^-/-^*BMDM’s were interferon-stimulated genes **(Figure 1F; Supplementary Figure 2C)**. Because of systemic induction of ISGs, again the top functions and diseases associated with *Irgm1* deficiency in brain and BMDMs were the antiviral response, systemic autoimmune syndrome (Systemic Lupus Erythematosus, Sjogren’s syndrome, Psoriasis), Rheumatic diseases, and other inflammatory disorders **(Figure 1G, Supplementary Figure 2D, Table S2)**. The RNA-seq data validation with qRT-PCR from brain tissues showed robust induction of key ISG’s (*Rig-I, Mda5, Ifi16, Irf7, Ifn-β, Stat1, Stat2, Isg15, Apobec3, Oas1, Mx1, and Mx2*) in *Irgm1^-/-^*mice (**Figure 1H**).

The class-I MHC restricted antigen presentation pathway is vital for processing and presentation of microbial (endogenous) and tumor antigens leading to antiviral/bacterial and anti-tumor response (Cresswell et al., 2005; Pamer and Cresswell, 1998). The expression of class 1 MHC genes is controlled by the interferon response (Coomans de Brachene et al., 2018; Keskinen et al., 1997; Zhou, 2009). Several of the genes integral to class-I MHC-mediated antigen processing and presentation pathways required for folding, assembly and peptide loading (HLA genes, immune-proteasome genes, B2M, and TAP1/2) (**Supplementary Figure 2E**) were upregulated in IRGM-depleted human and mice cells (**Supplementary Figure 2F**). Several of the complement pathway genes and the histone genes were upregulated in IRGM-depleted cells (**Supplementary Figure 2G, Table S1**). Both are known for their role in antimicrobial defense and the pathogenesis of systemic autoimmune diseases and are induced by interferon response (Byun et al., 2007; Chen et al., 2010; Chen et al., 2014; Mitchell et al., 1996; Silk et al., 2017). Several of the interferon-inducible TRIM proteins (TRIM5, 6, 12, 14, 20, 22 25, 29, 30, 34) that are known to play a significant role in innate immunity including inflammation and virus restriction (Ozato et al., 2008; van Gent et al., 2018) were significantly upregulated upon depleting IRGM (**Supplementary Figure 2H, Table S1**). Similarly, GBP’s that are the key effectors of the immune system against pathogens and are interferon responsive genes (Praefcke, 2018) were induced in IRGM-depleted cells (GBP1 and 4 in HT29 cell and GBP 2 to 10 in mice) (**Table S1**). The GO-based pathway analysis with genes that were downregulated in IRGM-depleted human or mice cells showed no immunity or inflammatory pathways suggesting that IRGM is a very specific suppressor of the inflammatory pathways (**Supplementary Figure 2I**).

Taken together, the transcriptome analysis in human cell and mice suggest that, (**1**) the IRGM-mediated regulation of immune systems and interferon response is systemic (not organ-specific), (**2**) the mouse Irgm1 (a 42 kDa protein) and human IRGM (a 21 kDa protein), although biochemically different, functionally are highly similar in regulation of inflammation especially in regulation of interferons responses, (**3**) IRGM is a master switch that suppresses the interferon responses under steady-state conditions and deficiency results in robust and systemic induction of type 1 IFN response.

### Constitutively activated PRR’s signaling pathways in IRGM depleted cells and mice

The mRNA expression of several cytoplasmic pattern recognition receptors (PRR’s), including *RIG-I, MDA5*, and *TLR3,* was significantly increased in *IRGM*-depleted mice and human cells **(Figure 2A)**. These PRR’s sense cytoplasmic DNA or dsRNA of self or pathogen origin and induces signaling events leading to the production of type I IFN’s, which then activate JAK-STAT signaling pathway for the production of ISG’s **(Supplementary Figure 3A)**. Through transcriptome analysis, it was explicit that the IRGM controls interferon response. However, what are the possible signaling pathways that induce the ISG’s in IRGM depleted cells was not evident. To understand this, we examined the expressions of proteins of DNA/RNA sensing and signaling pathways leading to ISG’s production (**Supplementary Figure 3A)** in IRGM human knockdown and mice knockout cells using western blotting experiments.

**Figure 2.**
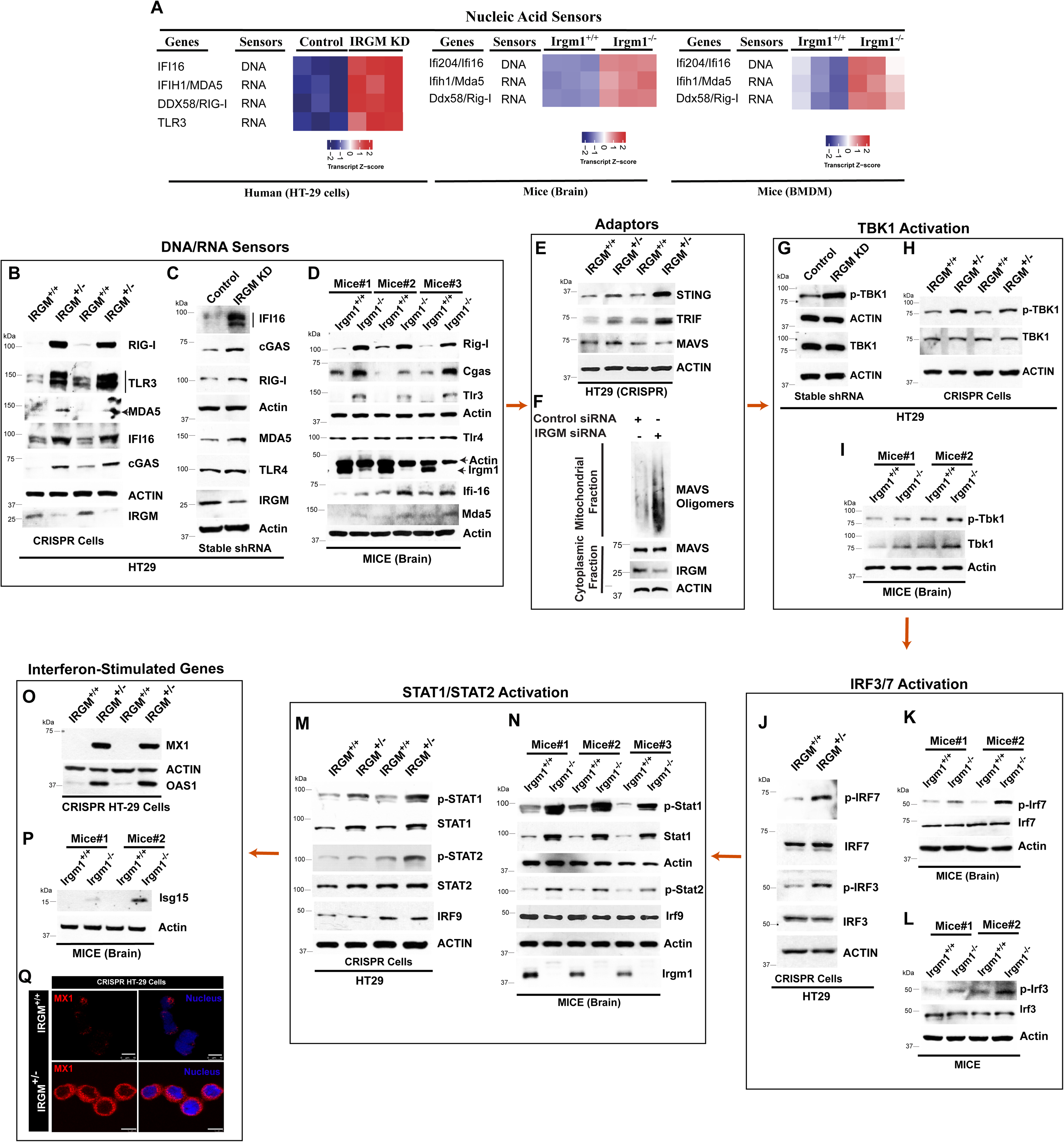
The nucleic acid-sensing and ISG production pathways are constitutively active in IRGM-depleted cells and mice. **(A)** Heat map of nucleic acid sensor proteins upregulated in IRGM KD HT29 cells and *Irgm1^-/-^* mice brain. **(B-D)** Western blot analysis to determine levels of nucleic acid sensor proteins with lysates of (**B**) HT29 control (**henceforth IRGM^+/+^**) and single allele CRISPR knockout IRGM cells (**henceforth IRGM^+/-^**) (2 biological repeats are shown, n=4), (**C**) HT29 cells stably expressing control shRNA or IRGM shRNA, (n=2) (**D**) *Irgm1^+/+^* and *Irgm1^-/-^* mice brain (n=3 mice). **(E)** Western blot analysis to determine levels of adaptor proteins in control and IRGM^+/-^ HT29 cells (2 biological repeats are shown, n=3). **(F)** SDD-AGE followed by western blot analysis with a mitochondrial fraction from control and IRGM siRNA knockdown cells. Western blot analysis with cytoplasmic fraction was also performed. (n=3) **(G-I)** Western blot analysis performed with lysates of (**G**) control and stable IRGM shRNA knockdown HT29 cells (**H**) IRGM^+/+^ and IRGM^+/-^ HT29 cells (2 biological repeats are shown, n=3), and (**I**) *Irgm1^+/+^* and *Irgm1^-/-^*mice brain to determine levels of TBK1 protein (n=2 mice). **(J-L)** Western blot analysis performed with lysates of (**J**) IRGM^+/+^ and IRGM^+/-^ HT29 cells (**K-L**) *Irgm1^+/+^* and *Irgm1^-/-^*mice brain to determine levels of IRF proteins (2 biological repeats or mice, n=3). **(M-N)** Western blot analysis performed with lysates of (**M**) IRGM^+/+^ and IRGM^+/-^ HT29 cells (2 biological repeats, n=3), (**N**) *Irgm1^+/+^*and *Irgm1^-/-^* mice brain to determine levels of STAT proteins (n=3 mice). **(O-P)** Western blot analysis performed with lysates of (**O**) IRGM^+/+^ and IRGM^+/-^ HT29 cells (2 biological repeats are shown, n=3) (**P**) *Irgm1^+/+^*and *Irgm1^-/-^* mice brain to determine levels of ISG proteins (n=2 mice). **(Q)** Representative confocal image of immunofluorescence assay performed with IRGM^+/+^ and IRGM^+/-^ HT29 cells stained with MX1 antibodies.

Even after several attempts, we were not able to generate and/or maintain complete CRISPR/Cas9 knockout of IRGM in THP1 or HT29 cells. The transfected cells were dying after a few days in culture. However, knockout of a single allele of IRGM was well tolerated in HT29 cells (Clone#7, henceforth **IRGM^+/-^**, **Supplementary Figure 3B**), which is used in several experiments in this study.

We observed increased protein expression of DNA and RNA sensor proteins RIG-I, TLR3, MDA5, IFI16, and cGAS in IRGM knockdown cells and *Irgm1^-/-^* mice (**Figure 2B-2D**). The TLR4 amount remained unchanged (**Figure 2C and 2D**). The cGAS was not induced at mRNA level in RNA-seq data, but at protein levels, an evident increase was observed. The adaptor proteins STING, MAVS, and TRIF transduce the signals from cGAS, RIG-I, and TLR3, respectively, leading to activation of TBK1 (**Supplementary Figure 3A**). The total amounts of STING and TRIF were higher in IRGM-depleted cells; however, MAVS levels were unchanged (**Figure 2E, Supplementary Figure 3C**). Although the total amount of MAVS was not increased, the MAVS aggregation, which is a hallmark of MAVS activation (Hou et al., 2011), was markedly induced in the absence of IRGM in SDD-AGE (Semi-Denaturating Detergent Agarose Gel Electrophoresis) assays (**Figure 2F**). To ascertain, we also performed immunofluorescence assays with IRGM depleted cells (**Supplementary Figure 3D and 3E**). The results clearly showed increased aggregation of MAVS in IRGM depleted mice BMDM’s and THP-1 cells (**Supplementary Figure 3D and 3E**). Also, these aggregates were co-localized over the mitochondria (**Supplementary Figure 3E)**. This data indicates that MAVS is activated in IRGM depleted cells and mice.

TBK 1 plays a central role in interferon response and serves as an integrator of multiple signals induced by nucleic acid sensors signaling cascades (cGAS, RIG-I, TLR3, and MDA5) leading to the activation of IRF3 and IRF7 transcription factors (**Supplementary Figure 3A**). TBK1 is activated by autophosphorylation at residue Ser172 (Shu et al., 2013). We observed increased phosphorylation of TBK1 in IRGM deficient human HT29 cell line and BMDM’s of *Irgm1^-/-^* mice (**Figure 2G-I)**. Although the total amount of TBK1 was unchanged in HT29, there was an increased expression of Tbk1 in BMDM’s of *Irgm1^-/-^*mice compared to controls. Activated TBK1 can increase phosphorylation of IRF3 and IRF7. Consistent with TBK1 activation in IRGM deficient cells, the activating phosphorylation (Ser396) of IRF3 and IRF7 (Ser477) was increased in IRGM/Irgm1 knockdown/knockout cells (**Figure 2J-2L**). However, the total amount of IRF3 and IRF7 remain unchanged.

When phosphorylated, IRF7 and IRF3 form homodimers or heterodimers translocate to the nucleus and induces the expression of the IFN genes (**Supplementary Figure 3A**). The IFN’s in autocrine or paracrine manner through the IFN receptors activate the JAK-STAT1/2 pathway (**Supplementary Figure 3A**). Next, we analyzed the status of the JAK-STAT signaling pathway by measuring the phosphorylation status of STAT1 and STAT2, the key events for the activation of this pathway. The total amount, as well as activating phosphorylation of STAT1 (Tyr701) and STAT2 (Tyr690), was substantially increased in IRGM knockout mice and knockdown cells (**Figure 2M and 2N, Supplementary Figure 3F**). The expression of IRF9 remained unchanged (**Figure 2M and 2N**). As a final step, we determined the protein expression of a few of the ISG’s. Heightened levels of MX1, OAS1, and Isg15 proteins were observed in IRGM deficient cells (**Figure 2O-2Q**). Altogether, the transcriptomic data followed by western blot analysis show that IRGM deficiency results in constitutive expression of several nucleic acid sensor proteins and activation of downstream interferon signaling pathways leading to increased JAK-STAT1/2 signaling resulting in enduring production of ISG’s.

### IRGM interacts and degrades nucleic acid sensor proteins to controls the aberrant activation of the interferon response

IRGM is an autophagy protein, and its deficiency leads to diminished autophagy flux in immune cells (Chauhan et al., 2015; Dong et al., 2015; Hansen et al., 2017; Kumar et al., 2018; Mehto et al., 2019; Singh et al., 2006). IRGM is known to interact and degrade several PRRs, including NOD1, NOD2, and NLRP3 (Chauhan et al., 2015; Mehto et al., 2019). The PRR’s are the threat sensor proteins, which commits the pathways for cytokines response. Hence, under steady-state conditions, maintaining their low expression is key to preserve the anti-inflammatory state of the cells. The total amount of several of the nucleic acid sensor proteins is controlled by autophagy-mediated degradation (Chen et al., 2016; Liu et al., 2016; Xian et al., 2019). We hypothesized that IRGM by mounting selective autophagy of nucleic acid sensor proteins restricts the type 1 IFN response under basal conditions. To validate this hypothesis, first, we examined whether IRGM interacts with the nucleic acid sensor proteins. In immunoprecipitation assays, a clear interaction between endogenous IRGM and endogenous RIG-I, cGAS, and TLR3 was observed **(Figure 3A**). However, no interaction was observed with MDA5, AIM2, TLR7, TLR4, and NLRC4 **(Figure 3A**), suggesting that IRGM specifically interacts with cGAS, RIG-I, and TLR3. We validated these interactions by performing co-immunoprecipitation assays (co-IP’s) in HEK293T cells with overexpressed proteins. GFP-IRGM clearly interacted with Flag-tagged cGAS, RIG-I, and TLR3 but not with AIM2 (**Figure 3B-3D, Supplementary Figure 4A**). The reverse co-IP’s were also performed to further validate these interactions (**Supplementary Figure 4B**).

**Figure 3.**
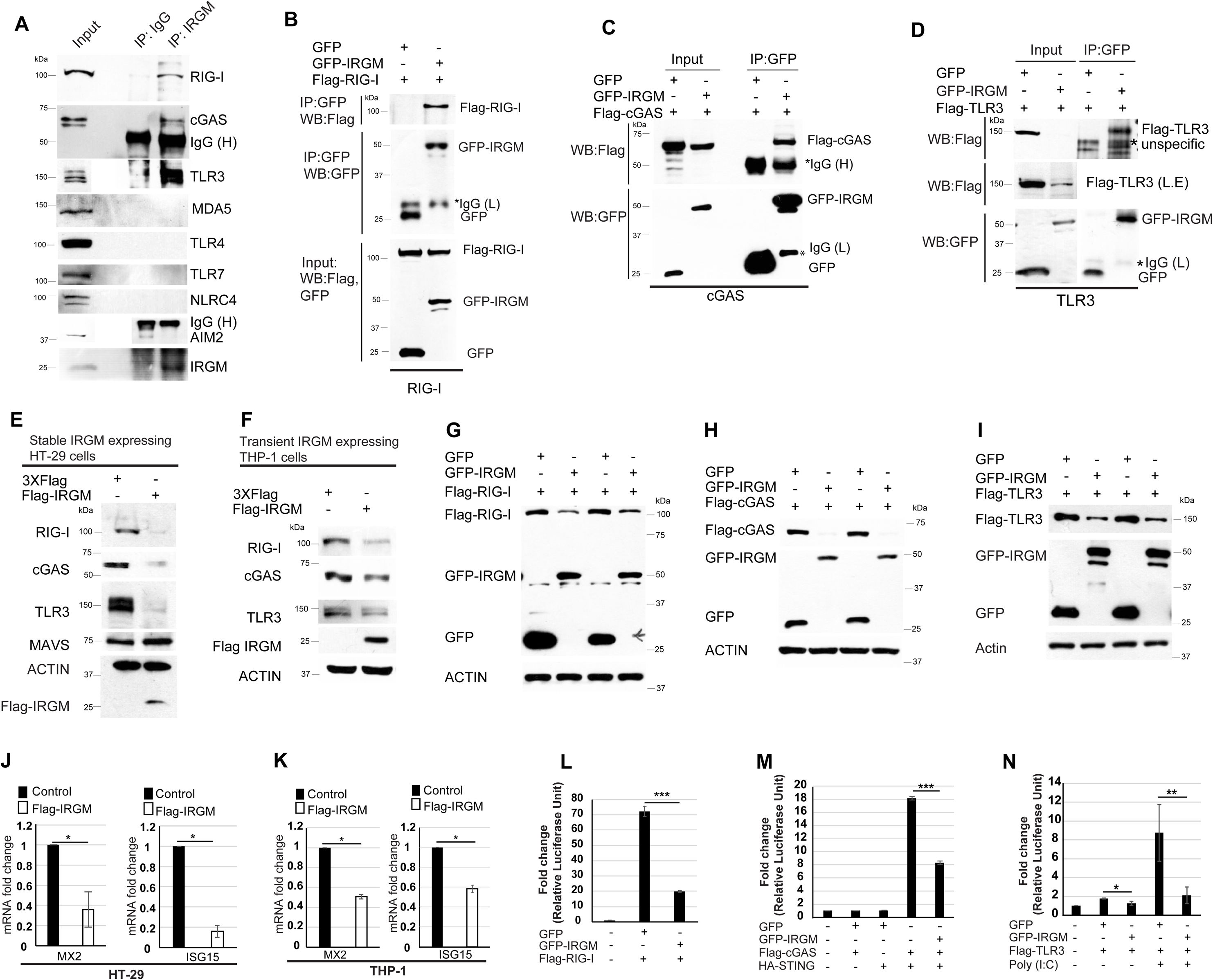
IRGM interacts and degrades cytoplasmic DNA and RNA sensor proteins to constrain IFN response. **(A)** Immunoprecipitation (IP) analysis of the interaction between endogenous IRGM and endogenous RIG-I, cGAS, MDA5, TLR3, TLR4, TLR7, NLRC4, and AIM2 in THP-1 cell lysates. IgG (H), IgG heavy chain. **(B-D)** Co-immunoprecipitation (Co-IP) analysis of the interaction between (**B**) GFP-IRGM and Flag-RIG-I, (**C**) GFP-IRGM, and Flag-cGAS, (**D**) GFP-IRGM and Flag-TLR3 in HEK293T cell lysates. L.E, Long exposure of input blot. IgG (L), IgG Light chain; IgG (H), IgG heavy chain. **(E)** Western blot analysis with the lysates of HT-29 cells stably transfected with control or Flag-IRGM plasmid and probed with the indicated antibodies. **(F)** Western blot analysis with lysates of THP-1 cell transiently transfected (4 h) with control or Flag-IRGM plasmid and probed with indicated antibodies. **(G-I)** Western blot analysis with lysates of HEK293T cells transiently expressing GFP or GFP-IRGM and (**G**) Flag-RIG-I or (**H**) Flag-cGAS or (**I**) or Flag-TLR3. (2 or 3 biological replicates are shown, n ≥ 3) **(J-K)** qRT-PCR analysis with RNA isolated from (**J**) control and Flag-IRGM stable HT-29 cells (**K**) THP-1 cells transiently transfected with control or Flag-IRGM plasmid. n=3, Mean ± SD, *P < 0.05, Student’s unpaired t-test. **(L-N)** Luciferase reporter assays in HEK293T cells transiently transfected with ISRE reporter plasmid and other plasmids as indicated. Mean ± SD, n = 3, **P < 0.005, ***P < 0.0005, Student’s unpaired t-test.

Next, we scrutinized whether IRGM can degrade the cytoplasmic sensor proteins to which it interacts. The endogenous levels of RIG-I, cGAS, and TLR3 (but not of TLR4 and MAVS) were reduced by stable (HT-29 cells) or transient (THP1 cells) overexpression (for 4 hrs) of Flag-IRGM (**Figure 3E and 3F)**. Previously, we have shown that IRGM does not affect the expression levels of NLRC4 and AIM2 (Mehto et al., 2019). Since cGAS, RIG-I, and TLR3 expression is controlled by IFN response, the reduction of endogenous levels of these proteins could be an indirect effect of IRGM-mediated suppression of IFN response. To rule out this possibility, we overexpressed both IRGM and the sensor proteins using CMV promoter-driven ORF’s in HEK293T cells. The results clearly show that IRGM overexpression can reduce the total amount of RIG-I, cGAS, and TLR3 (**Figure 3G-I)** but not of AIM2, MAVS, and TLR4 **(Supplementary Figure 4A and 4C),** suggesting that IRGM is directly involved in the degradation of RIG-I, cGAS, and TLR3 sensor proteins. This phenomenon is further validated by the overexpression of Flag-tagged IRGM (**Supplementary Figure 4D)**.

Consistent with these results, the overexpression of IRGM suppressed the levels of the sentinel ISG genes, *MX2,* and *ISG15* (**Figure 3J and 3K)**. Furthermore, in ISRE (Interferon-stimulated response element) luciferase reporter assays, the overexpression of IRGM reduced the RIG-I, cGAS/STING, and TLR3 induced ISRE-driven promoter transcription (**Figure 3L-3N)**. Overall, the data suggest that IRGM interacts and degrades RIG-I, cGAS, and TLR3 to keep type I IFN response under-check.

### IRGM mediates p62-dependent autophagic degradation of nucleic acid sensors to restrains the activation of the interferon response

Using autophagy and proteasome inhibitors, we next determined the process utilized by IRGM to degrade these proteins. IRGM mediated degradation of endogenous RIG-I, cGAS, and TLR3 were abrogated by autophagy/lysosomal inhibitors; Bafilomycin A1 (BafA1) and chloroquine (**Figure 4A and 4B).** The proteasomal inhibitor, MG132, was not able to block the IRGM-mediated degradation of endogenous RIG-I and TLR3, whereas it diminished the degradation of cGAS (**Supplementary Figure 4E).** Similarly, in overexpression experiments, the GFP-IRGM mediated degradation of Flag-RIG-I and Flag-TLR3 was abrogated by autophagy inhibitors but not by MG132 (**Figure 4C-4E)** whereas cGAS degradation was reduced by both BafA1 and MG132 (**Figure 4D)**. This data suggest that IRGM majorly invokes autophagic degradation of the RIG-I and TLR3, however cGAS expression is controlled by both autophagic and proteasomal degradation. If autophagy is the key process employed by IRGM to degrade the nucleic acid sensors, then IRGM-mediated suppression of ISG’s should be rescued by autophagy inhibitors. Indeed, inhibition of autophagy by BafA1 significantly de-repressed the IRGM-mediated inhibition of expression of ISG’s (**Figure 4F).**

**Figure 4.**
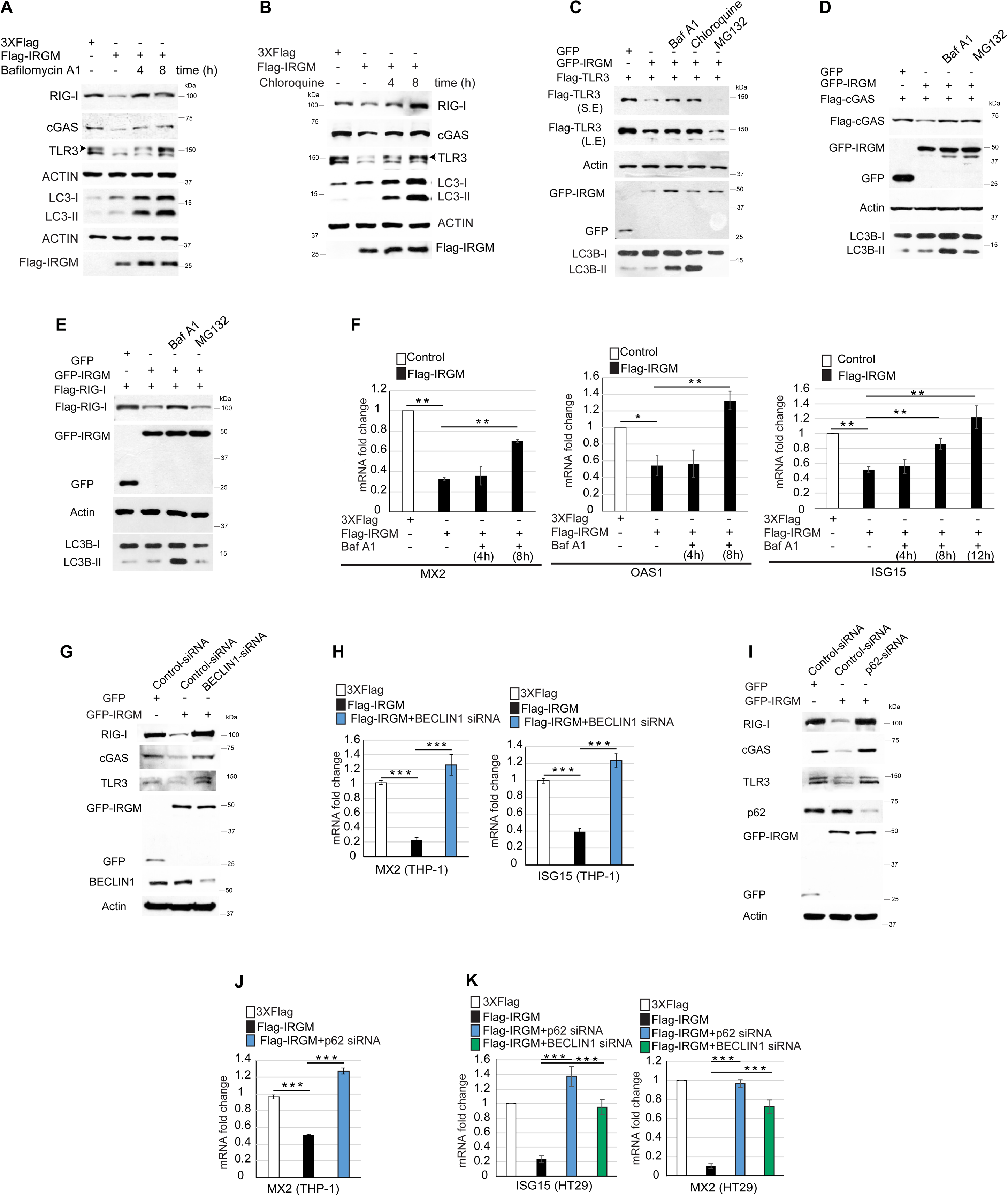
IRGM mediates p62-dependent selective autophagy of nucleic acid sensors. **(A)** Western blotting analysis with lysates of control and untreated or Bafilomycin A1 (300 nM; 4 h and 8 h) treated Flag-IRGM expressing HT29 stable cell lines. (n=3) **(B)** Western blotting analysis with lysates of control and untreated or chloroquine (50 µM; 4 h and 8 h) treated Flag-IRGM expressing HT29 stable cell lines. (n = 3) **(C-E)** HEK293T cells expressing GFP or GFP-IRGM and (**C**) Flag-TLR3, (**D**) Flag-cGAS, and (E) Flag-RIG-I were treated with Bafilomycin A1 (300 nM, 8h) or chloroquine (50 μM, 8h) or MG132 (20 μM, 8h) and cell lysates were subjected to western blot analysis with the indicated antibodies. (n=3) **(F)** qRT-PCR analysis with RNA isolated from untreated or Bafilomycin A1 (300 nM; 4 h, 8 h, and 12 h) treated control or Flag-IRGM expressing HT-29 stable cells as indicated. Mean ± SD, n = 3, *P < 0.05, **P < 0.005 Student’s unpaired t-test. **(G)** The cell lysates of control or BECLIN1 siRNA transfected THP-1 cells expressing GFP or GFP-IRGM were subjected to immunoblotting with indicated antibodies. **(H)** qRT-PCR analysis with RNA isolated from control or BECLIN1 siRNA transfected THP-1 cells expressing 3X-Flag epitope or Flag-IRGM as indicated. Mean ± SD, n = 3, ***P < 0.0005, Student’s unpaired t-test. (**I**) The cell lysates of control or p62 siRNA transfected THP-1 cells expressing GFP or GFP-IRGM (as indicate) were subjected to immunoblotting with indicated antibodies. **(J)** A qRT-PCR analysis with RNA isolated from control or p62 siRNA transfected THP-1 cells expressing 3X-Flag epitope or Flag-IRGM as indicated. Mean ± SD, n = 3, ***P < 0.0005, Student’s unpaired t-test. **(K)** qRT-PCR analysis with RNA isolated from control or p62 siRNA or BECLIN1 siRNA transfected HT29 cells expressing 3X-Flag epitope or Flag-IRGM as indicated. Mean ± SD, n = 3, ***P < 0.0005, Student’s unpaired t-test.

IRGM facilitates the autophagic degradation of proteins. However itself, it is not degraded by autophagy (Kumar et al., 2018; Mehto et al., 2019). This is not surprising as several of the core autophagy proteins such as ULK1, ATG16L1, ATG12 (Haller et al., 2014; Nazio et al., 2016; Scrivo et al., 2019) and endolysosomal trafficking proteins such as RAB7A (Mohapatra et al., 2019), which facilitates autophagic degradation of cargo proteins but themselves are not degraded by the autophagy.

Above, we used chemical inhibitors to show that IRGM induces autophagy to degrade sensor proteins and control type 1 IFN response. Next, we used genetic method to validate this finding. BECLIN1 is an essential autophagy protein required for the initiation of canonical autophagy. The siRNA mediated knockdown of BECLIN1 in IRGM overexpressing cells abolished the IRGM mediated degradation of nucleic acid sensor proteins (**Figure 4G).** Further, siRNA mediated depletion of BECLIN1 IRGM overexpressing cells fully restored the expression of ISG’s in two different cell lines (**Figure 4H and 4K)** suggesting that IRGM mounts canonical BECLIN-1 dependent autophagy as a key mechanism to maintaining low levels of nucleic acid sensor proteins and keeping the type I IFN response under-check.

The autophagy adaptor proteins recognize cargoes for selective autophagic degradation (Kim et al., 2016; Svenning and Johansen, 2013). We screened the interaction between IRGM and several established autophagy adaptor proteins, including Optineurin, TAX1BP1, p62, NDP52, and NBR1, to identify the adaptor protein/s utilized by IRGM to mediate selective autophagic degradation of sensor proteins. IRGM interacted strongly with p62, mildly with TAX1BP1, and no interaction was observed with Optineurin, NDP52, and NBR1 (**Supplementary Figure 4F and 4G)**. Several of the key PRR’s (including cGAS and RIG-I) and immune proteins (including STING and TRIF) are shown to be degraded by p62-mediated autophagy (Chen et al., 2016; Liu et al., 2016; Prabakaran et al., 2018; Samie et al., 2018; Xian et al., 2019). From our IRGM-p62 interaction data and the literature, we envisaged that p62 could be the key adaptor protein employed by IRGM for selective autophagic degradation of nucleic acid sensor proteins. Indeed, the depletion of p62 in IRGM overexpressing cells abrogated the IRGM-mediated degradation of RIG-I, cGAS, and TLR3 (**Figure 4I)** and also rescued the IRGM-mediated downregulation of ISG’s (**Figure 4J and 4K)**. Altogether, the data suggest that IRGM complexes with p62 for selective autophagic degradation of RIG-I, cGAS, and TLR3 leading to a controlled type I IFN response under basal conditions.

### IRGM deficiency results in defective mitochondrial flux and increased mtROS

IRGM deficiency results in increased expression of nucleic acid sensor proteins and activation of interferon signaling pathways (**Figure 1 and 2**). However, it was not clear to us, what keeps the sensors in ON state, and fuels the persistent activation of these pathways. Next, we set to determine the cell intrinsic stimuli (DAMPs) that fuels the enduring IFNs signaling in *Irgm1^-/-^* mice. In the absence of external stimuli, the endogenous DAMPs from dysfunctional mitochondria are the major triggers for the type 1 IFN response (Angajala et al., 2018; West et al., 2015; Zhang et al., 2010). We and others have shown that depletion of IRGM in human cells or mice results in defective autophagy (Dong et al., 2015; He et al., 2012; Mehto et al., 2019) (**Supplementary Figure 5A and 5B**) and this can result in impaired mitophagy and accumulation of defunct mitochondria. Indeed, we observed a significantly higher number of fused, swollen, and clumped mitochondria in *Irgm1^-/-^* BMDMs and IRGM siRNA knockdown cells (**Figure 5A-5D)**. Next, we scrutinized the status of mitophagy in these cells. An E3-ubiquitin ligase, Parkin mediates mitophagy downstream of ubiquitin kinase, PINK1 (PTEN-induced kinase 1) (Jin and Youle, 2012; Narendra et al., 2008; Vives-Bauza et al., 2010). The PINK1 and PARKIN themselves are degraded by mitophagy, and a defect in productive mitophagy will result in their accumulation over defective mitochondria. In immunofluorescence assays, we observed substantially increased localization of endogenous PARKIN and PINK1 over the clumped mitochondria in *Irgm1^-/-^* BMDMs compared to control cells (**Figure 5E and 5F, Supplementary Figure 5C**). This phenomenon was more pronounced when YFP-Parkin was overexpressed in BMDMs (**Supplementary Figure 5D**). In agreement, in western blot analysis, we observed an accumulation of both PARKIN and PINK1 in IRGM depleted human and mice macrophages compared to the control cells (**Figure 5G and 5H**). Further, in the presence of autophagy inhibitor Bafilomycin A1, the accumulation of PARKIN and PINK1 is increased in control cells but not in IRGM depleted cells (**Figure 5I and 5J**) suggesting a block of mitophagy flux in IRGM/Irgm1 knock out and knockdown cells. The data suggest that mitophagy flux is impaired in IRGM depleted cells leading to an accumulation of abnormal clumped mitochondria in these cells.

**Figure 5.**
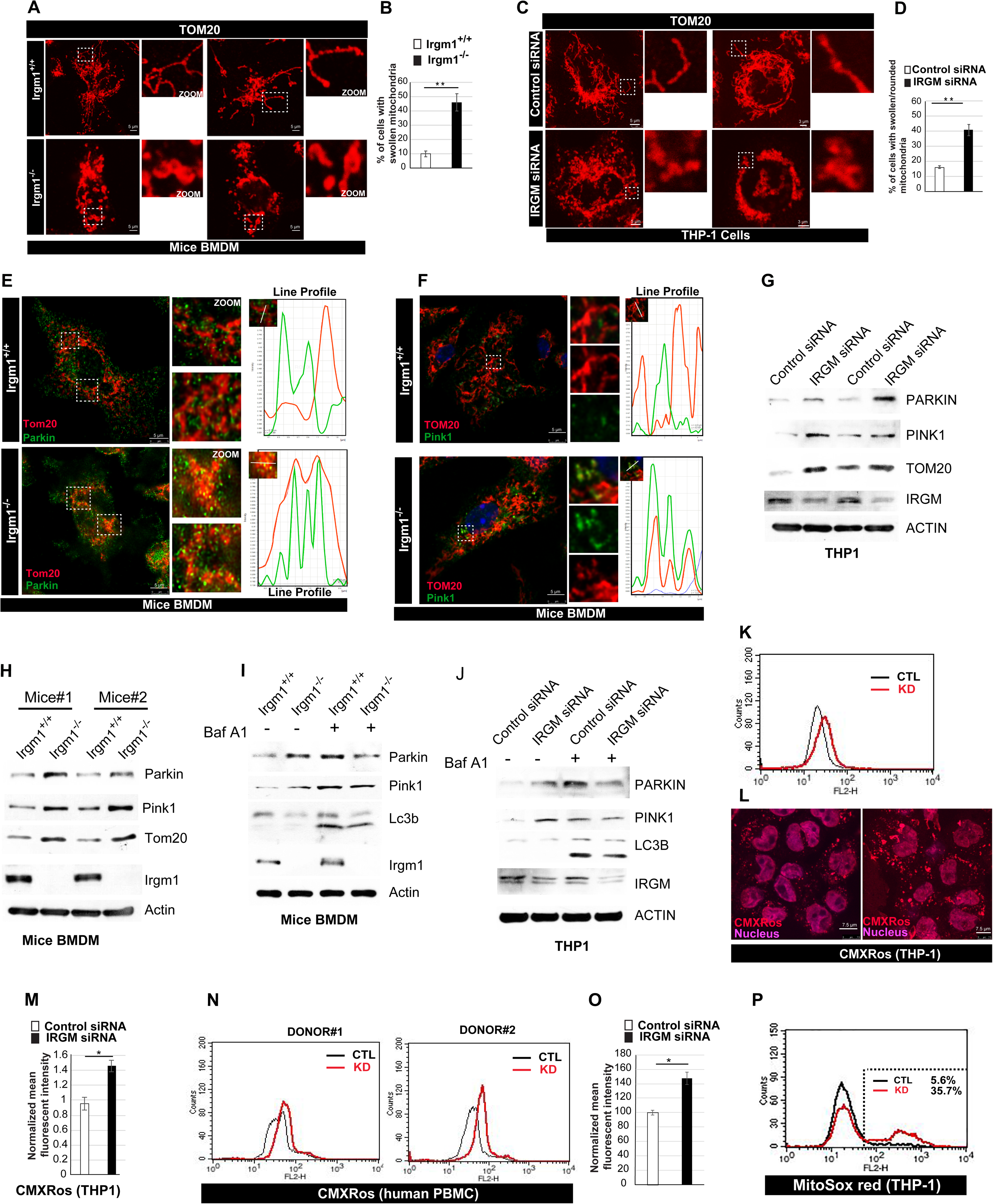
IRGM depletion results in impaired mitophagy and increased mtROS. **(A)** Representative confocal images of *Irgm1^+/+^* and *Irgm1^-/-^* mice BMDMs processed for IF analysis with Tom20 antibody. **(B)** The graph depicts the percentage of cells with swollen/rounded mitochondria in *Irgm1^+/+^* and *Irgm1^-/-^*mice BMDMs. Mean ± SD, n = 3, **P < 0.005, Student’s unpaired t-test. **(C)** Representative confocal images of control siRNA and IRGM siRNA transfected THP-1 cells processed for IF analysis with TOM20 antibody. **(D)** The graph depicts the percentage of cells with swollen or rounded mitochondria in control siRNA and IRGM siRNA transfected THP-1 cells. Mean ± SD, n = 3, **P < 0.005, Student’s unpaired t-test. **(E)** Representative confocal (STED) images of *Irgm1^+/+^* and *Irgm1^-/-^* mice BMDMs processed for IF analysis with Tom20 and Parkin antibodies. Line profile: Co-localization analysis using line intensity profiles. **(F)** Representative confocal (STED) images of *Irgm1^+/+^* and *Irgm1^-/-^* mice BMDMs processed for IF analysis with Tom20 and Pink1 antibodies. Line profile: Co-localization analysis using line intensity profiles. **(G)** Western blot analysis of mitochondrial fraction of control siRNA and IRGM siRNA transfected THP-1 cells, probed with indicated antibodies (2 biological replicates shown, n = 3). **(H)** Western blot analysis of mitochondrial fraction of *Irgm1^+/+^* and *Irgm1^-/-^*mice BMDM cells, probed with indicated antibodies (n=2 mice). **(I)** Western blot analysis of mitochondrial fraction of *Irgm1^+/+^*and *Irgm1^-/-^* mice BMDM cells untreated or treated with Bafilomycin A1 (300 nM, 3 h), probed with indicated antibodies. **(J)** Western blot analysis of mitochondrial fraction of control siRNA and IRGM siRNA transfected THP-1 cells untreated or treated with Bafilomycin A1 (300 nM, 3 h), probed with indicated antibodies. **(K)** Flow cytometry analysis of control siRNA and IRGM siRNA transfected THP-1 cells stained with CMXRos red dye (10 nM, 30 mins). **(L)** Representative confocal images of control and IRGM siRNA transfected THP-1 cells stained with CMXRos red dye. **(M)** The Graph depicts the normalized mean fluorescent intensity of control and IRGM knockdown THP-1 cells stained with CMXRos. Mean ± SD, n = 3, *P < 0.05, Student’s unpaired t-test. **(N)** Flow cytometry analysis of control siRNA and IRGM siRNA transfected human PBMC’s from two donor’s stained with CMXRos red dye (10 nM, 30 mins). **(O)** The graph depicts the normalized mean fluorescent intensity of control and IRGM knockdown human PBMC’s stained with CMXRos red dye. Mean ± SD, n = 2, *P < 0.05, Student’s unpaired t-test. **(P)** Representative flow cytometry analysis of control and IRGM siRNA transfected THP-1 cells stained with MitoSox red dye (1μM, 20 min). (n=4). The microscopic scales are as depicted in the images

Next, we used Mitotracker CMXRos^TM^ dye to measure mitochondrial membrane potential. IRGM knockdown THP-1 cells and PBMC’s from healthy human donors showed increased CMXRos staining compared to the control cells in flow cytometry analysis and immunofluorescence (**Figure 5K-5O**) suggesting an increased polarization of mitochondria in IRGM deficient cells compared to the control cells. To further verify this finding, we performed JC-1 mitochondrial staining, whose accumulation is dependent only on membrane potential but not on size, shape, and density. At low mitochondrial potential, JC-1 is predominantly a monomer and yields green fluorescence, whereas, at the higher mitochondrial potential, it forms aggregates and yield red fluorescence. The IRGM deficiency leads to increased red fluorescence (red to green ratio), suggesting a hyperpolarized state of the mitochondria (**Supplementary Figure 5E and 5F)**. This data is consistent with previous observations where overexpression of IRGM leads to mitochondrial fission and depolarization (Singh et al., 2010), and here we observed fused and hyperpolarized mitochondria in IRGM deficient cells. The hyperpolarized state of mitochondria is often associated with the production of mitochondrial reactive oxygen species (mtROS) leading to apoptosis or necrosis and also autoimmune diseases (Angajala et al., 2018; Chen et al., 2017; Galloway and Yoon, 2012; Gergely et al., 2002a; Gergely et al., 2002b; Perl et al., 2004). Indeed, the mitochondrial superoxide and cellular total ROS levels, as measured by MitoSOX™ and CellRox™ dyes, were elevated in IRGM-depleted THP-1 cells, mice BMDM’s and human PBMC’s (**Figure 5P, Supplementary figure 5G and 5H).** Taken together, the data suggest that cellular depletion of IRGM results in defective mitophagy flux, accumulation of fused hyperpolarized mitochondria, and increased oxidative stress.

### The cytosol of IRGM deficient cells is soiled with DAMPs that fuels IFN response

The endogenous dsDNA needs to be strictly compartmentalized within the nucleus or mitochondria, and any leakage of dsDNA into cytosol results into strong immune stimulation leading to inflammation and autoimmunity (Crow and Rehwinkel, 2009; Roers et al., 2016). The increase in cellular oxidative stress results in mitochondrial and genomic instability leading to the release of mtDNA and nuclear DNA (micronuclei) into the cytosol (Gehrke et al., 2013). The mtDNA or micronuclei induces strong interferon response via cGAS-STING-IRF3 axis leading to autoimmune and auto-inflammatory conditions (Ablasser et al., 2014; Mackenzie et al., 2017; Pisetsky, 2012; West et al., 2015; White et al., 2014). Since IRGM depletion results in the accumulation of defective mitochondria, and increased oxidative stress, we examined the status of dsDNA species in the cytosol. Immunofluorescence assays in both human and mice cells showed increased mtDNA nucleoids in the cytosol (**Figure 6A and 6B)**. The micronuclei (both lamin B1 bound or unbound), and extracellular DNA was also substantially increased in IRGM-depleted cells (**Figure 6C-6F, Supplementary Figure 5I, and 5J)**. Extracellular DNA is another potent DAMP, which is released from necrotic cells and can gain access to the cytoplasm of surviving cells to induce inflammation (Pisetsky, 2012). To examine whether cytosolic DNA is responsible for augmented type I IFN response in IRGM knockdown cell, we electroporated DNase I enzyme in these cells to deplete cytosolic DNA and then performed qRT-PCR to measure ISG’s expression. No cell toxicity was observed in cells due to DNase 1 electroporation. The DNase 1 treatment (5 µg) for 1 hour significantly reduced the elevated expression of ISG’s in IRGM knockdown cells (**Figure 6G)**. This data suggest that cytosolic DNA is a DAMP that provokes type I IFN signaling in IRGM-depleted cells.

**Figure 6.**
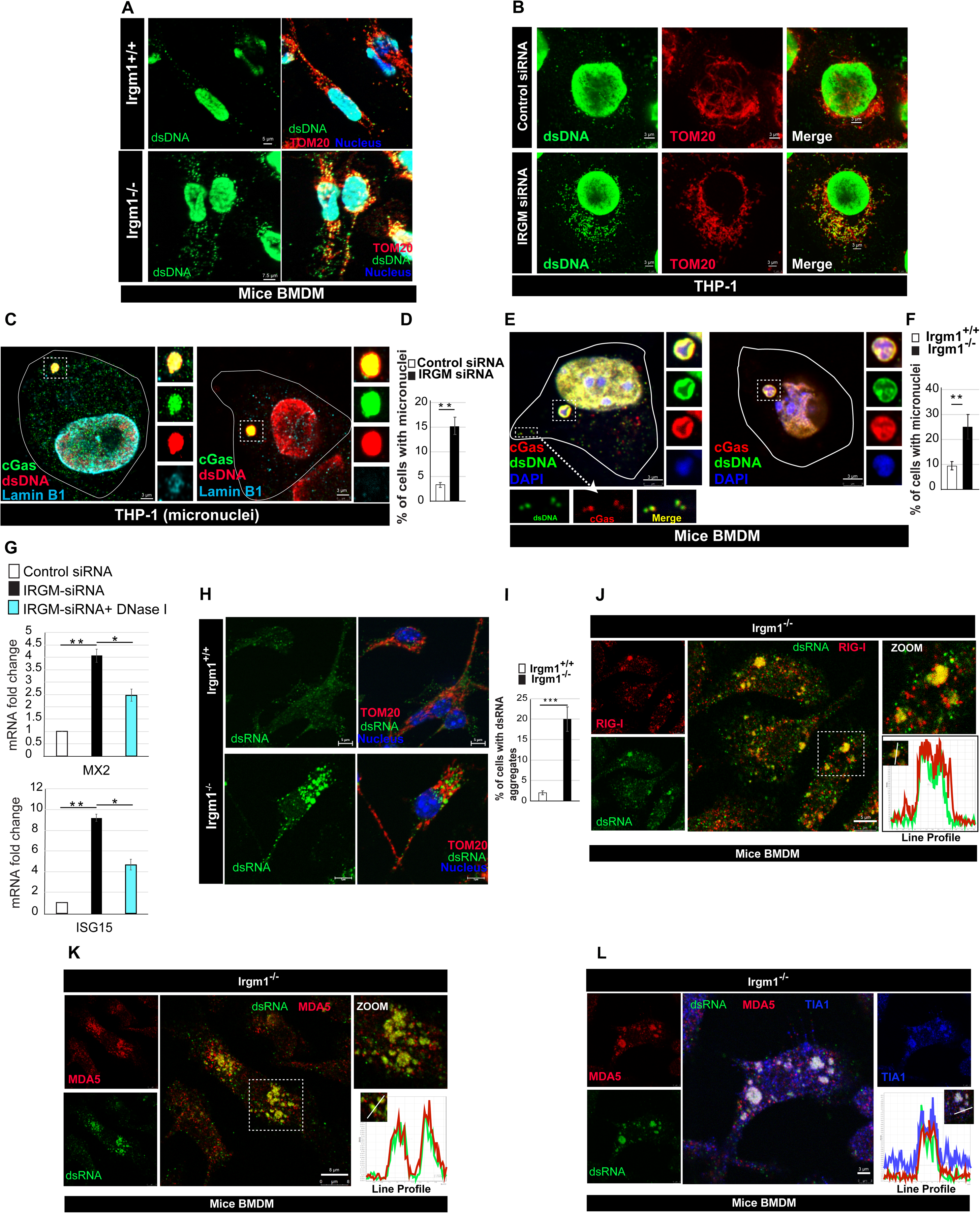
IRGM depleted cells are soiled with mtDAMPs that induces IFN response. **(A)** Representative confocal images of *Irgm1^+/+^* and *Irgm1^-/-^* mice BMDMs processed for IF analysis with Tom20 (red) and dsDNA (green) antibody. **(B)** Representative confocal images of siRNA and IRGM siRNA transfected THP-1 cells processed for IF analysis with TOM20 (red) and dsDNA (green) antibodies. **(C)** Representative STED microscopy images of control and IRGM knockdown THP-1 cells processed for IF analysis with cGAS (green), dsDNA (red) and Lamin B1 (cyan) antibodies. **(D)** Graph depicts the percentage of cells with micronuclei in control siRNA and IRGM siRNA knockdown THP-1 cells. Mean ± SD, n = 3, **p < 0.005, Student’s unpaired t-test. **(E-F)** Representative STED microscopy images of *Irgm1^-/-^* mice BMDMs **(E)** transfected with mCherry-cGAS and immunostained with dsDNA (green) antibody. **(F)** The graph depicts the percentage of cells with micronuclei in *Irgm1^+/+^*and *Irgm1^-/-^* mice BMDMs. Mean ± SD, n = 3, **p < 0.005, Student’s unpaired t-test. **(G)** The qRT-PCR analysis with RNA isolated from control and IRGM siRNA knockdown THP-1 cells electroporated with DNase I (5 μg, 1hr) as indicated. Mean ± SD, n = 3, *p < 0.05, **p < 0.005, Student’s unpaired t-test. **(H-I)** Representative confocal images of *Irgm1^+/+^*and *Irgm1^-/-^* mice BMDMs processed for IF analysis with (H) dsRNA (green) and Tom20 (red) antibodies. **(I**)The graph depicts the percentage of *Irgm1^+/+^* and *Irgm1^-/-^* mice BMDMs with dsRNA aggregates. Mean ± SD, n = 3, ***p < 0.0005, Student’s unpaired t-test. **(J)** Representative confocal images of *Irgm1^-/-^*mice BMDMs processed for IF analysis with RIG-I (red) and dsRNA (green) antibodies. Line profile: Co-localization analysis using line intensity profiles. **(K-L)** Representative confocal images of *Irgm1^-/-^* mice BMDMs processed for IF analysis with (**K**) Mda5 (red) and dsRNA (green) antibodies or (**L**) Mda5 (red), dsRNA (green) and Tia1 (blue) antibodies. Line profile: Co-localization analysis using line intensity profiles. The microscopic scales are as depicted in the images

The viral and mitochondrial dsRNA can trigger the RIG-I/MDA5-MAVS-IRF3 signaling pathway leading to enhanced interferon production (Dhir et al., 2018; Linder and Hornung, 2018; Reikine et al., 2014). We performed immunofluorescence assays with extensively used dsRNA-specific J2 antibody to explore the possibility of the presence of dsRNA in the cytosol of IRGM-depleted cells. A low level of endogenous dsRNA, which partly co-localizes with mitochondria, was observed in control cells (**Figure 6H and 6I, Supplementary Figure 6A)**. In a very surprising observation, a significantly higher number of *Irgm1^-/-^*BMDM’s showed an abundant number of dsRNA structures in the cytosol (**Figure 6H and 6I, Supplementary Figure 6A)**. Endogenous RIG-I and MDA5 showed complete co-localization with these dsRNA structures (**Figure 6J and 6K, Supplementary Figure 6B)**. Since we found that IRGM-deprived cells are under constitutive oxidative stress, we tested whether these dsRNA structures are the stress granules. Indeed, the stress granules marker, TIA1, completely co-localized with the dsRNA structures suggesting that they are the stress granules (**Figure 6L)**. We electroporated dsRNA specific RNase (RNase III) and RNA-DNA hybrid-specific RNase (RNase H) in the IRGM knockdown cells to deplete cytosolic dsRNA and then performed qRT-PCR to measure ISG’s expression. Contrary to DNase 1 exposure, the treatment of RNase resulted in increased cell toxicity, so the treatment was performed only for one hour (no cell death at this time point). Nevertheless, the heightened ISG’s expression in IRGM depleted human and mice BMDM’s was significantly reduced upon RNase treatment **(Supplementary Figure 6C and 6D).** Our attempt to electroporate both DNAse I and RNase’s together is failed due to cytotoxicity. From the data presented here, we conclude that increased DNA and dsRNA in the cytosol of IRGM deficient cells trigger enduring type 1 IFN signaling and response.

### Both cGAS-STING and RIG-I/MDA5-MAVS signaling contributes to enduring type 1 IFN response in IRGM-depleted cells

The cytosolic DNA sensor cGAS was found to be perfectly co-localized with micronuclei (**Figure 6C and 6E)** and also with cytosolic nucleoid (**Figure 6E)** in the IRGM knockout cells. This indicates that the cGAS-STING axis could account for heightened type I IFN response and augmented ISG’s production in IRGM-depleted cells. Indeed, the siRNA knockdown of cGAS and STING in IRGM-depleted HT29, THP-1, and mice BMDM cells significantly diminished the increased expression of IFN-β and ISG’s (MX2 and ISG15) (**Figure 7A-7E)**.

**Figure 7.**
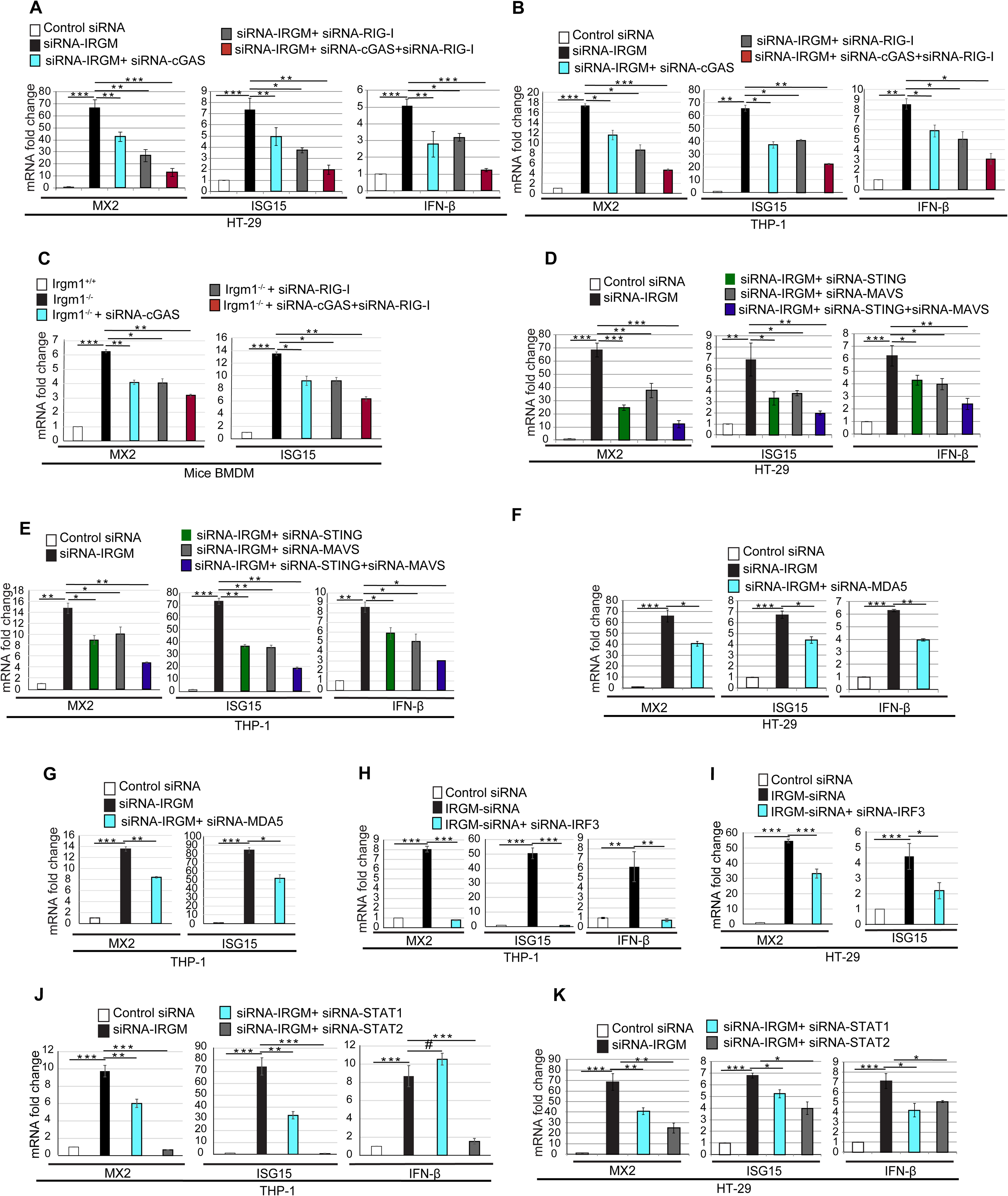
The cGAS-STING and RIG-I-MAVS signaling synergistically activate IFN response in IRGM-depleted cells. **(A-B)** qRT-PCR analysis with total RNA isolated from (**A**) HT-29 (**B**) THP-1 cells transfected with control siRNA or IRGM siRNA or doubly transfected with IRGM siRNA and cGAS siRNA or IRGM siRNA and RIG-I siRNA or transfected with three siRNA’s as indicated. Mean ± SD, n = 3, *p < 0.05, **p < 0.005, ***p < 0.0005, Student’s unpaired t-test. **(C)** qRT-PCR analysis with total RNA isolated from *Irgm1^+/+^* and *Irgm1^-/-^* mice BMDMs transfected with control siRNA or cGAS siRNA or RIG-I siRNA as indicated. Mean ± SD, n = 3, *p < 0.05, **p < 0.005, ***p < 0.0005, Student’s unpaired t-test. **(D-E)** qRT-PCR analysis with total RNA isolated from (**D**) HT-29 (**E**) THP-1 cells transfected with control siRNA or IRGM siRNA or doubly transfected with IRGM siRNA and STING siRNA or IRGM siRNA and MAVS siRNA or transfected with all the three siRNA’s as indicated. Mean ± SD, n = 3, *p < 0.05, **p < 0.005, ***p < 0.0005, Student’s unpaired t-test. **(F-G)** qRT-PCR analysis with total RNA isolated from (**F**) HT-29 (**G**) THP-1 cells transfected with control siRNA or IRGM siRNA or doubly transfected with IRGM siRNA and MDA5 siRNA as indicated. Mean ± SD, n = 3, *p < 0.05, **p < 0.005, ***p < 0.0005, Student’s unpaired t-test. **(H-I)** qRT-PCR analysis with total RNA isolated from (**H**) THP-1 (**I**) HT-29 cells transfected with control siRNA or IRGM siRNA or doubly transfected with IRGM siRNA and IRF3 siRNA as indicated. Mean ± SD, n = 3, *p < 0.05, **p < 0.005, ***p < 0.0005, Student’s unpaired t-test. **(J-K)** qRT-PCR analysis with total RNA isolated from (**J**) THP-1 (**K**) HT-29 cells transfected with control siRNA or IRGM siRNA or doubly transfected with IRGM siRNA and STAT1 siRNA or IRGM siRNA and STAT2 siRNA as indicated. Mean ± SD, n = 3, *p < 0.05, **p < 0.005, ***p < 0.0005, Student’s unpaired t-test

Upon viral infection, RIG-I localizes over the stress granules to enhance type 1 IFN response (Kuniyoshi et al., 2014; Onomoto et al., 2012). The presence of RIG-I/MDA5 in the atypical dsRNA stress granules in IRGM knock out BMDMs (**Figure 6J and 6K, Supplementary Figure 6B)** could be important for the activation of RIG-I/MDA5-mediated IFN signaling. Next, using siRNA, we depleted RIG-I, MDA5, MAVS, and IRF3 in IRGM knockdown cells (**Supplementary Figure 6E-J**) to explore whether RIG-I/MDA5-MAVS-IRF3 axis contributes to elevated type I IFN levels. Indeed, the knockdown of RIG-I, MDA5, MAVS, and IRF3 in IRGM-depleted human and mice cells, abated the heightened type I interferon response as scored by expression of IFN-β and ISG’s (**Figure 7A-7I)**.

Since both DNA (cGAS-STING) and RNA (RIG-I-MAVS) sensing axis appears to contribute to augmented IFN response in IRGM deficient cells, we tested whether depleting the sensors/adaptors of both pathways simultaneously may have a synergistic effect. Indeed, double knockdown of cGAS and RIG-I or STING and MAVS in IRGM-depleted HT29 and THP-1 cells resulted in a greater decrease of type 1 IFN response as compared to individual knockdowns (**Figure 7A-7E**) suggesting that both DNA and RNA sensing pathways are activated in IRGM deficient cells leading to augmented and sustained type I IFN response.

STAT1 and STAT2 are the transcription factors of the JAK-STAT pathway, which binds to ISRE (Interferon stimulated response element) to induce transcription of ISGs. In THP-1 cells, the depletion of STAT2 in IRGM knockdown cells completely abolished the increased expression of ISGs (**Figure 7J)**, whereas STAT1 depletion has a marginal effect (**Figure 7J)** suggesting that STAT2 is more important for the increased expression of ISGs in IRGM-depleted THP-1 cells. However, in HT-29 cells, both STAT1 and STAT2 appears to be equally important for increased expression of ISG’s (**Figure 7K).**

It appears that increased ROS in IRGM-depleted cells plays a role in triggering the events, eventually inducing the ISG’s. We tested whether depletion of ROS by N-acetyl-L-cysteine (NAC) affects the level of ISG’s in IRGM knockdown cells. Indeed, exposure of IRGM-depleted cells with NAC for just two hours significantly dampens the induced expression of ISG’s (**Supplementary Figure 6K),** suggesting that certainly, ROS plays an important role in eliciting the events finally leading to heightened type I IFN response in IRGM knockdown cells.

Collectively, the data suggest that the events triggered by reduced mitophagy flux in IRGM depleted cells culminating in the production of DAMPs stimulates DNA/RNA sensing-signaling axis to produce IFNs, which then activates JAK-STAT signaling pathway for ISG’s production.

### Depletion of IRGM establishes an intrinsic antiviral state in human cells and mice

The interferon responsive ISG’s are the major defense arsenal against the viruses (Crosse et al., 2018; Dixit and Kagan, 2013). The restriction factors, which are known to inhibit virus attachment, membrane fusion, uncoating, transcription, translation, assembly, release, and reinfection (**Supplementary Figure 7A**) were highly elevated at RNA (**Figure 8A, Supplementary Figure 7B-7D**) and protein levels (**Figure 8B)** in IRGM-depleted cells indicating a strong antiviral state of the cells. Indeed, a potent inhibition of replication of CHIKV (ssRNA virus, Alphavirus, Togaviridae), HSV-1 (dsDNA virus, Simplexvirus, Herpesviridae) and JEV (ssRNA virus, Flavivirus, Flaviviridae) as measured by qRT-PCR (with primers specific to viruses) was observed in IRGM knockdown cells (**Figure 8C-8E**). The plaque assay is used as the gold standard for virus quantification as it can directly measure the infectious viral particle titer. In plaque assays, there was a strong reduction in total infectious CHIKV and HSV-1 virus particles produced by IRGM knockdown cells compared to the wild type cells (**Figure 8F and 8G**).

**Figure 8.**
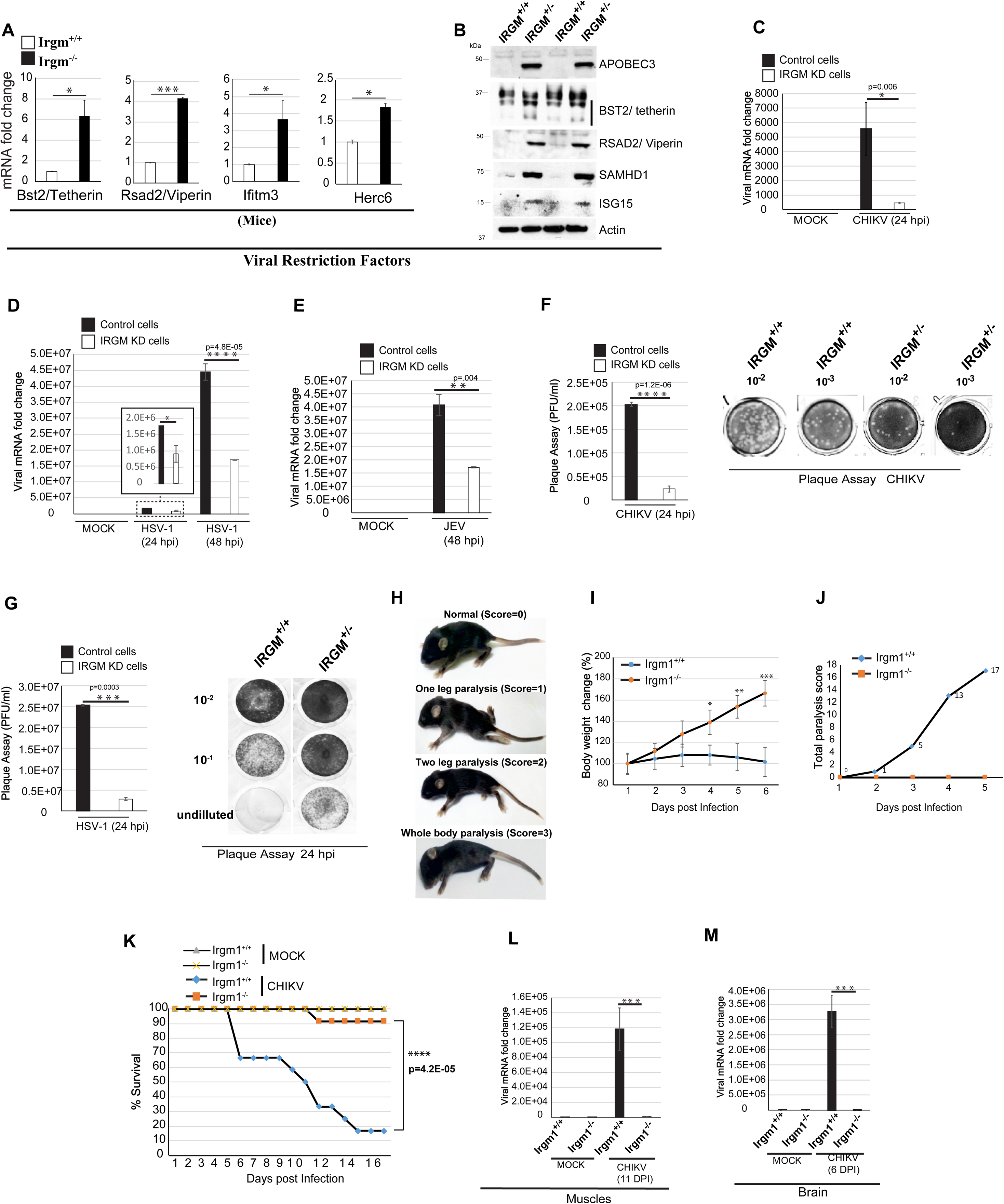
The depletion of IRGM stimulates the cell-intrinsic antiviral state. **(A-B)** qRT-PCR analysis with indicated viral restriction factor genes from the total RNA of **(A)** *Irgm1^+/+^* and *Irgm1^-/-^* mice brain and (**B**) Immunoblot analysis performed with lysates of IRGM^+/+^ and IRGM^+/-^ HT29 cells with indicated antibodies. Mean ± SD, n = 3, *p < 0.05, **p < 0.005, ***p < 0.0005, Student’s unpaired t-test. **(C)** Total RNA was isolated from CHIKV (MOI=5) infected control and IRGM knockdown HT29 cells subjected to qRT-PCR with CHIKV specific primers to quantitate total viral load. Mean ± SE, n = 3, *p < 0.05, Student’s unpaired t-test. **(D)** Total RNA was isolated from HSV-1 (MOI=1) infected control and IRGM knockdown HT29 cells subjected to qRT-PCR with HSV-1 specific primers to quantitate total viral load. Mean ± SE, n = 3, *p < 0.05, ****p < 0.00005, Student’s unpaired t-test. **(E)** Total RNA was isolated from JEV (MOI =1) infected control and IRGM knockdown HT29 cells subjected to qRT-PCR with JEV specific primers to quantitate total viral load. Mean ± SE, n =3, **p < 0.005, Student’s unpaired t-test. **(F)** Left panel, quantification of CHIKV plaque assays (plaque-forming unit/ml) in Vero cells performed from culture supernatant (24 hpi) of HT29 control and IRGM knockdown cell. n=3, Mean ± SE, ****p < 0.00005, Student’s unpaired t-test. Right panel, representative images of plaque assays. **(G)** Left panel, quantification of HSV-1 plaque assays (plaque forming unit/ml) in Vero cells performed from culture supernatant (24 hpi) of HT29 control and IRGM knockdown cell. n=3, Mean ± SE, ***p < 0.0005, Student’s unpaired t-test. Right panel, representative images of plaque assays performed with different dilutions of virus-containing culture supernatant. **(H)** The stages of paralysis in C57BL/6 mice post CHIKV infection (6 dpi) with scoring according to the diseases conditions. **(I)** Graph depicts the percentage change in body weight of CHIKV infected *Irgm1^+/+^* and *Irgm1^-/-^* neonates (n=12, 3 different experiments) for duration as indicated. Mean ± SD, *P < 0.05, **P < 0.005, ***P < 0.0005, Student’s unpaired t-test. **(J)** Graph depicts total paralysis scores of CHIKV infected *Irgm1^+/+^* and *Irgm1^-/-^*neonates (n=12, 3 different experiments) until 5 days post infection **(K)** Graph depicts the percentage survival of mock and CHIKV infected *Irgm1^+/+^* and *Irgm1^-/-^* neonates (n=12, 3 different experiments) during the course of infection. **(L)** Total RNA isolated from muscles of mock and CHIKV infected (11 dpi) *Irgm1^+/+^* and *Irgm1^-/-^* mice and was subjected to qRT PCR for quantitation of viral load (Mean ± SD, n = 3, ***P < 0.0005, Student’s unpaired t-test). **(M)** Total RNA isolated from brains of mock and CHIKV infected (6 dpi) *Irgm1^+/+,^* and *Irgm1^-/-^* mice and was subjected to qRT PCR for quantitation of viral load (Mean ± SD, n = 3, ***P < 0.0005, Student’s unpaired t-test). The total RNA used for qRT-PCR with brain is four times more than muscles

To validate this important finding *in vivo*, we employed a CHIKV infection neonatal mice model (Couderc et al., 2008). In this model, the young age and the inefficient type I IFN signaling were found to be the reasons for severe disease outcomes (Couderc et al., 2008). Intradermal injection of 10^7^ PFU’s (plaque-forming units) to 6 days old wild type C57BL/6 neonates can induce disease symptoms. Typically, a mild paralysis in one leg is observed on day 3^rd^ post-infection (pi), severe leg paralysis is observed 6-7 day pi (dpi) followed by the death of C57BL/6 mice by 12-15 dpi (**Figure 8H**). All the CHIKV infected wild type neonate mice displayed no significant increase in body weight until 6 dpi, followed by a reduction in body weight until they died (**Figure 8I**). However, a constant increase in body weight of *Irgm1^-/-^* neonates were observed until the termination of the experiments (**Figure 8I**). All of the twelve infected wild type neonate mice developed typical symptoms of paralysis by 5-6^th^ dpi (**Figure 8H and 8J**), and eleven of them died by the 15^th^ dpi (**Figure 8K**). In striking contrast, there were no visible symptoms of paralysis in all the twelve *Irgm1^-/-^* neonate mice by 5 dpi (**Figure 8K**). Later, two mice developed mild paralysis, and one died on 12 dpi (**Figure 8K**). None of the other ten *Irgm1^-/-^*neonate mice showed any disease symptoms until the termination of the experiments (**Figure 8K, data not shown**). This data shows that a very strong intrinsic antiviral state exists in neonates of *Irgm1^-/-^* mice, which can easily clear up the high dose of CHIKV virus infection. We measured the viral load in muscles and brain of wild type and *Irgm1* knockout mice. There was a robust increase in viral load in both muscles and brains of wild type mice (**Figure 8L and 8M**). Consistent with symptoms, the *Irgm1^-/-^*mice displayed a very significant reduction in viral load (almost negligible viral load) in muscles and brain (**Figure 8L and 8M**). Taken together, the *in vitro* and the *in vivo* results demonstrate that IRGM could be a potent therapeutic target for inducing broad-spectrum antiviral immunity.

## Discussion

This study reveals that IRGM is a master switch that controls the levels of type I IFN response by controlling the activation of DNA/RNA sensing inflammatory pathways. A prominent role of IRGM in suppressing inflammation presented here or previously (Chauhan et al., 2015; Mehto et al., 2019), justify why IRGM protein, which was dead for 20 million years of evolution might have revived back in ancestors of humans (Bekpen et al., 2009).

### IRGM is a molecular link between autophagy and inflammation

IRGM/Irgm1 is an important autophagy protein, which recruits Syntaxin17 over the autophagosomes and play significant role in autophagosome-lysosome fusion (Chauhan et al., 2015; Dong et al., 2015; Hansen et al., 2017; Kumar et al., 2018; Mehto et al., 2019; Singh et al., 2006; Singh et al., 2010). IRGM interacts and stabilizes core autophagy proteins, ULK1, ATG16L1, and BECLIN1 (Chauhan et al., 2015). IRGM acts as a scaffold protein and provides a platform for interaction of NOD2 and autophagy proteins to mount antimicrobial autophagy and to control NFĸB activation in the immune cells (Chauhan et al., 2015). Utilizing similar autophagy machinery, IRGM mediates degradation of NLRP3 and ASC to control IL-1β production in the inflammatory bowel disease mouse model (Mehto et al., 2019). In this study, we found that IRGM selectively interacts and mediates autophagic degradation of cGAS, RIG-I, and TLR3 to control interferon response. Taken together, these studies suggest that IRGM, because of its capability to interact with both autophagy proteins and inflammatory signaling proteins, can act as a molecular link between autophagy and inflammation. Specifically, it can utilize the autophagy machinery to degrade inflammatory proteins to maintain immune homeostasis.

### Targeting IRGM for anti-viral therapeutics

A robust antiviral state was observed in IRGM-depleted cells and *Irgm1^-/-^* mice. Several mechanisms might have contributed to such powerful broad-spectrum antiviral response, including potent induction of RNA/DNA sensing and type I IFN pathways, upregulation of almost all viral restriction factors, the presence of upregulated MHC-1 antigen presentation/processing pathway, complement pathway, and persistent presence of stress granules. In the CHIKV neonate infection mice model, we observed that *Irgm1^-/-^* neonates (just six days old) were almost completely protected from viral pathogenesis, suggesting a very strong innate immunity to the viruses in these mice. We cautiously speculate that therapeutic targeting of IRGM could provide new broad-spectrum antivirals.

### Loss of IRGM invokes a vicious cycle of events that endorse cellular Inflammation

The SNP’s and deletion polymorphisms in promoter region of *IRGM* and SNP in exonic region, which changes the binding capacity of microRNA-196 to *IRGM* was shown to be strongly associated with susceptibility to Crohn’s diseases and other autoimmune diseases (Brest et al., 2011; McCarroll et al., 2008; Xia et al., 2017; Yao et al., 2018). Both of these types of genetic changes reduce IRGM expression (Brest et al., 2011; McCarroll et al., 2008), suggesting that decreased expression of IRGM enhances the susceptibility to autoimmune and auto-inflammatory diseases. Our, this and a previous study (Mehto et al., 2019), shows that depletion of IRGM triggers a vicious cycle of inflammation leading to increase susceptibility to inflammatory diseases. We found that in the absence of IRGM, cGAS-STING, RIG-I-MAVS, and NLRP3-ASC inflammatory pathways get activated, leading to the production of strong inflammatory cytokines, including IL-1β and type 1 IFN’s. The secretion of these cytokines could induce inflammatory cell death (Mehto et al., 2019), resulting in the release of DAMP’s, which in the feedback loop further fuels the inflammatory signaling pathways to produce more cytokines. We envisage that this vicious cycle of inflammation is triggered in patients of autoinflammatory diseases carrying mutations in the *IRGM* gene. Further studies are needed and will be taken up in human patients to demonstrate this hypothesis. Nevertheless, this study defines the mechanism by which IRGM is a master regulator of IFN response and highlights IRGM as a strong potential target for new therapeutic interventions against viral and inflammatory diseases.

## Acknowledgments

This work is funded by the Wellcome Trust/Department of Biotechnology (DBT) India Alliance (IA/I/15/2/502071) fellowship to Santosh Chauhan. Punit Prasad is supported by Ramalingaswami Re-entry fellowship of the Department of Biotechnology (BT/HRD/35/02/2006). Soma Chattopadhyay is funded by the Department of Science and Technology (DST), New Delhi, India, vide-grant no EMR/2016/000854. We acknowledge the technical assistance of Bhabani Sahoo (Microscopy facility), Kshitish Rout, and Paritosh Nath (FACS facility). We gratefully acknowledge the support of the Institute of Life Sciences central facilities funded by Department of Biotechnology (India). We gratefully acknowledge Dr. M. M. Parida, Dr. A. Basu, Dr. Roger Everett for providing CHIKV strain, JEV strain HSV-1 KOS strain, respectively.

## Author Contributions

S.C. secured funding, conceived the project, designed the experiments, and wrote the manuscript. K.K.J, S.M., P.N., and N.R.C, performed the majority of experiments. Swati.C, R.S, T.K.N, S.P.K., S.K.D., K.D, P.K.S., K.C.M, S.D, and A.D performed the experiments. Swati.C, P.P, and Soma.C provided critical inputs for experiments and edited the manuscript.

## Conflict of Interest

The authors declare that they have no conflict of interests.

## Supplementary Figure Legends

**Supplementary Figure 1.**
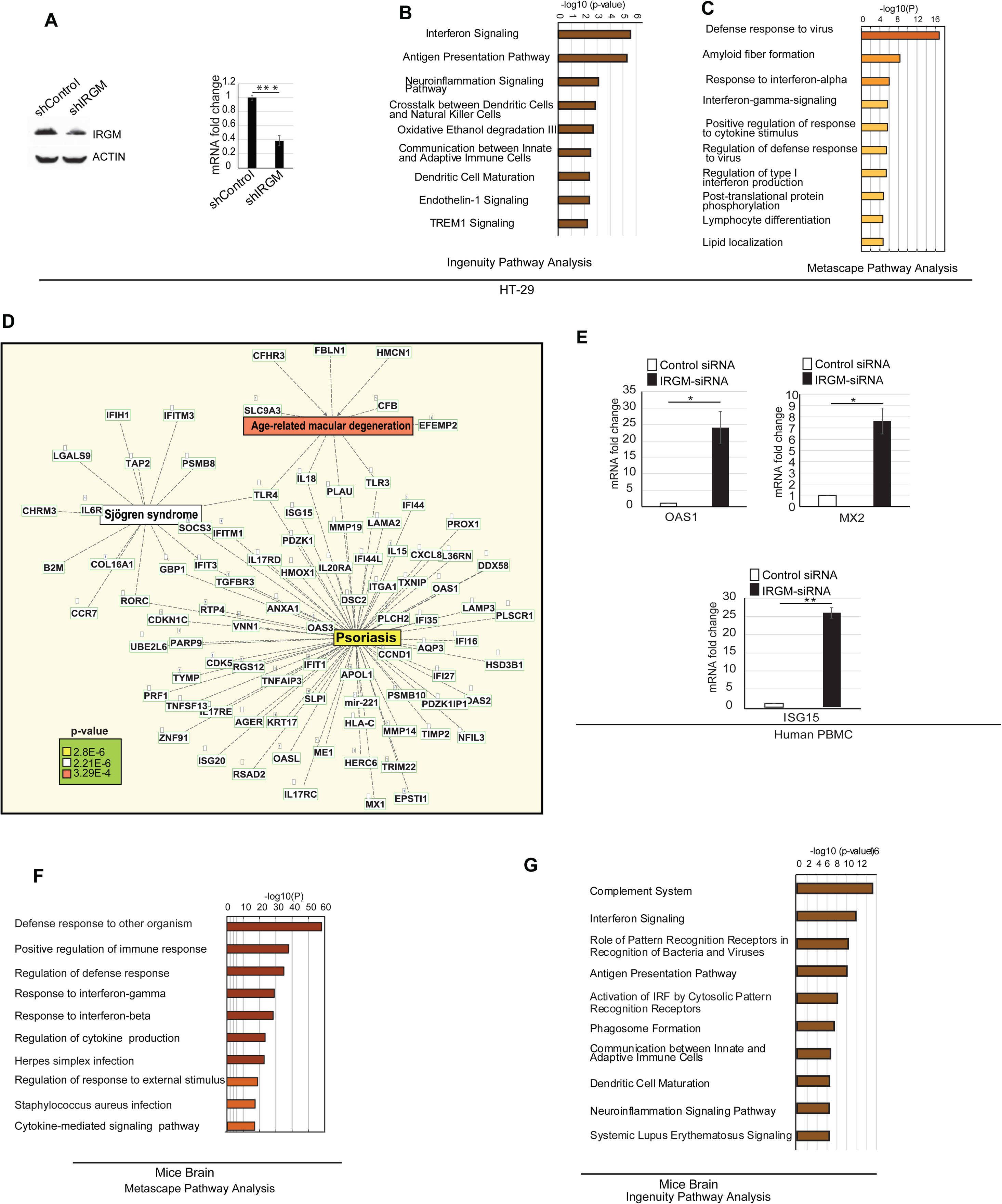
IRGM is a negative regulator of inflammatory signaling pathways. **(A)** Top panel, IRGM knockdown efficiency (around 50%) analyzed by western blot using cell lysate of HT29 stably expressing control shRNA or IRGM shRNA. Right panel, IRGM knockdown efficiency was analyzed using qRT-PCR from the same cells. (**B, C**) The bar graphs represent highly enriched 10 biological pathways upregulated in gene ontology (GO) based (**B**) metascape pathway analysis (**C**) Ingenuity pathway analysis using sets of genes induced (1.5 fold, p<0.05, n=3) in IRGM shRNA knockdown HT29 cells compared to control shRNA cells. **(D)** Network pathway analysis using IPA. The molecular network of genes connected with diseases associated with genes (1.5 fold, p<0.05) upregulated in *Irgm1^-/-^* mice brain. The complete list is documented in supplementary table S2. **(E)** qRT-PCR validation of RNA-seq data in control siRNA and IRGM siRNA transfected human PBMC cells. Mean ± SD, n = 3, *P < 0.05, **P < 0.005, Student’s unpaired t-test. (**F, G**) The bar graphs represent top pathways upregulated in GO-based (**F**) Metascape pathway analysis (G) Ingenuity pathway analysis using sets of genes induced (1.5 fold, p<0.05, n=3) in the brain of *Irgm1^-/-^*knockout mice compared to *Irgm1^+/+^* wild type mice.

**Supplementary Figure 2.**
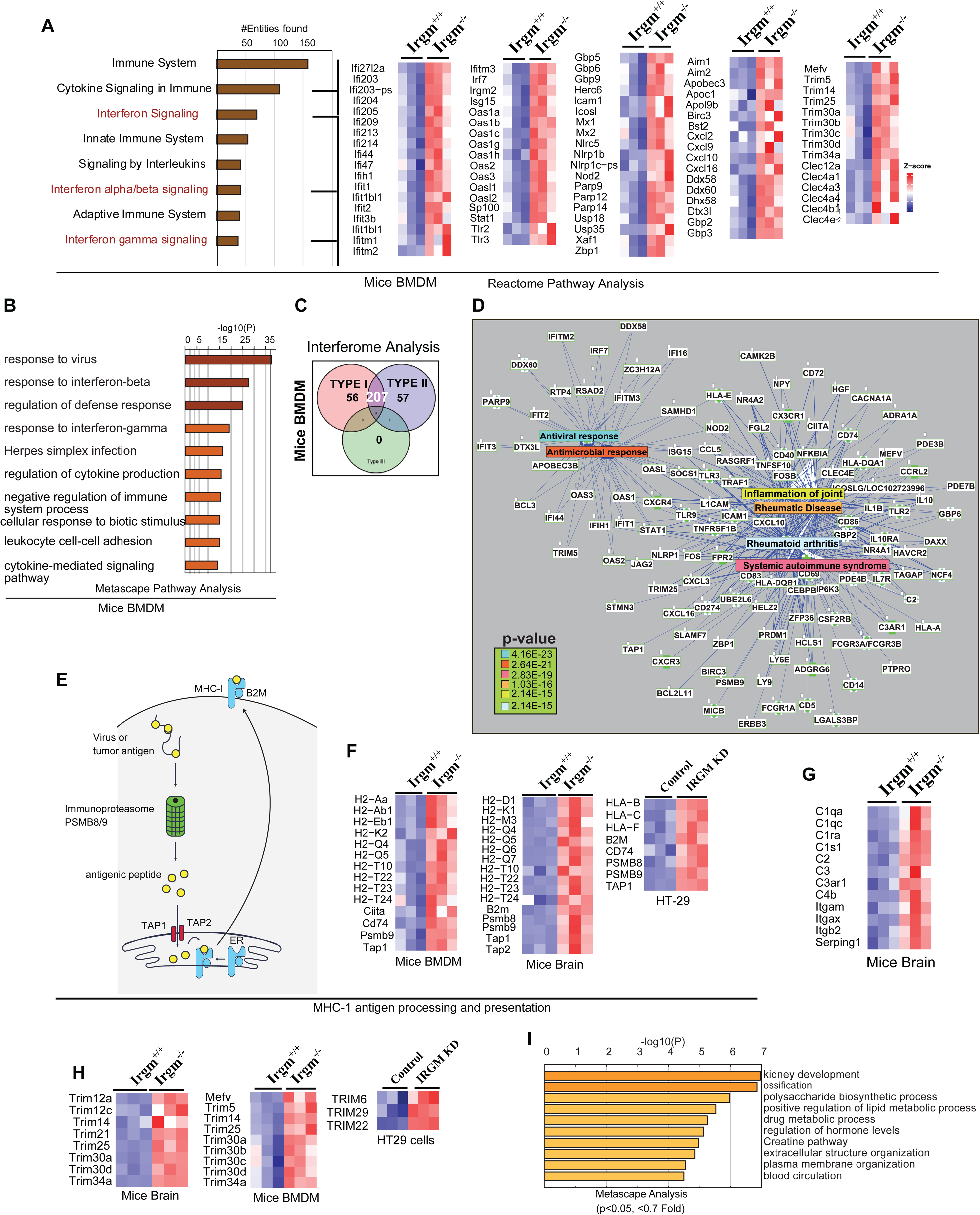
IRGM is a negative regulator of type-1 IFN signaling pathways in mice BMDM’s. (**A**). Bar graph represents top 10 canonical biological pathways upregulated in GO based reactome pathway analysis using sets of genes induced (1.5 fold, p<0.05, n=3) in *Irgm1^-/-^* mice BMDM’s compared to *Irgm1^+/+^*mice BMDM’s. Heatmaps were generated for sentinel interferon-regulated genes (three biological replicates). (**B**) Bar graph represent top 10 canonical biological pathways upregulated in GO-based metascape pathway analysis using sets of genes induced (1.5 fold, p<0.05, n=3) in *Irgm1* knockout mice BMDM’s compared to wild type mice BMDM’s. (**C**). Interferome database analysis with sets of genes induced (1.5 fold, p<0.05, n=3) in mice BMDM of *Irgm1^-/-^* mice compared to *Irgm1^+/+^*wild type mice. The Venn diagram depicts the total number of upregulated type I and type II IFN-regulated genes in *Irgm1^-/-^* mice BMDM’s. (**D**) Network pathway analysis using IPA. The molecular network of genes connected with diseases/function associated with genes (1.5 fold, p<0.05, RNA-seq) upregulated in *Irgm1* knockout mice BMDM’s. The complete list is documented in supplementary table S2. (**E-F**) Left panel, (E) diagrammatic representation of MHC-1 antigen processing and presentation pathway. (F) Right panels, heatmaps of the genes of this pathway differentially expressed in *Irgm1^+/+^* and *Irgm1^-/-^*mice BMDM’s and brain and also control and IRGM knockdown HT29 cells. **(G)** Heatmaps showing genes of complement system pathway upregulated (1.5 fold, p<0.05, n=3) in the brain of *Irgm1^-/-^* mice compared to *Irgm1^+/+^* mice. **(H)** Heatmaps of the TRIM family genes upregulated (1.5 fold, p<0.05, n=3) in *Irgm1^-/-^* mice BMDM’s and brain and also IRGM knockdown HT29 cells. **(I)** Bar graph represents top 10 canonical biological pathways upregulated in GO-based metascape pathway analysis using a set of genes repressed (≤ 0.7 fold, p<0.05) in *Irgm1* knockout mice BMDM’s compared to wild type mice BMDM’s.

**Supplementary Figure 3.**
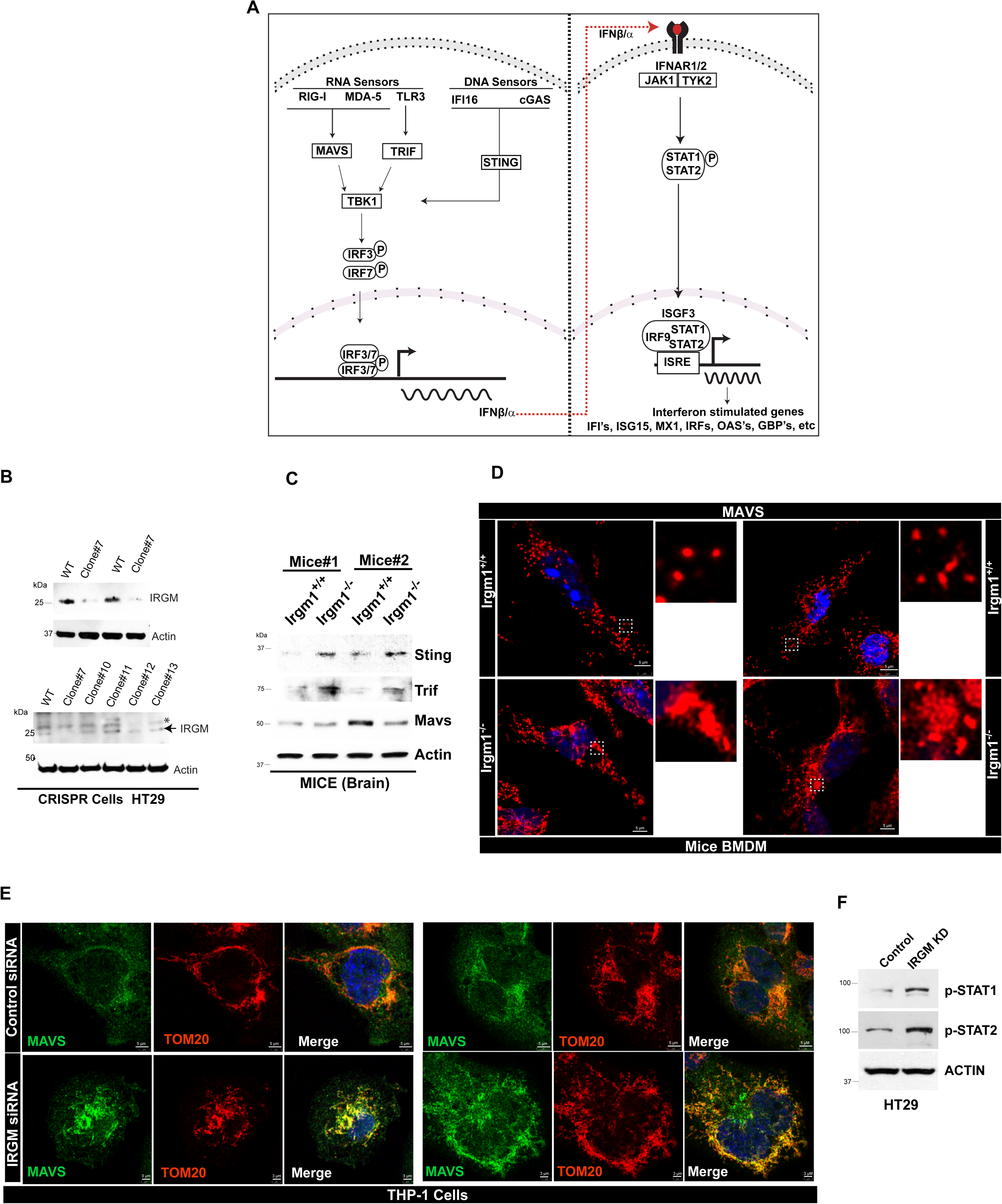
The nucleic acid sensing and ISG production pathways are constitutively active in IRGM-depleted cells. **(A)** The graphical representation of the pathways associated with type I IFN and ISG production. **(B)** Western blot analysis with lysates of wild type and a few CRISPR-Cas9 clones of HT29 cells. Clone #7 was identified to be singe allele knockout for IRGM.* Non-specific band. **(C)** Western blot analysis of adaptor proteins in *Irgm1^+/+^*and *Irgm1^-/-^* mice brain lysates probed with indicated antibodies. **(D)** Representative confocal images of *Irgm1^+/+^* and *Irgm1^-/-^* mice BMDMs immunostained with MAVS (red) antibody. **(E)** Representative confocal images of control siRNA and IRGM siRNA transfected THP-1 cells immunostained with MAVS (green) and TOM20 (red) antibodies. **(F)** Western blot analysis to assess activation of p-STAT1 and p-STAT2 in HT29 cells stably expressing control shRNA and IRGM shRNA.

**Supplementary Figure 4.**
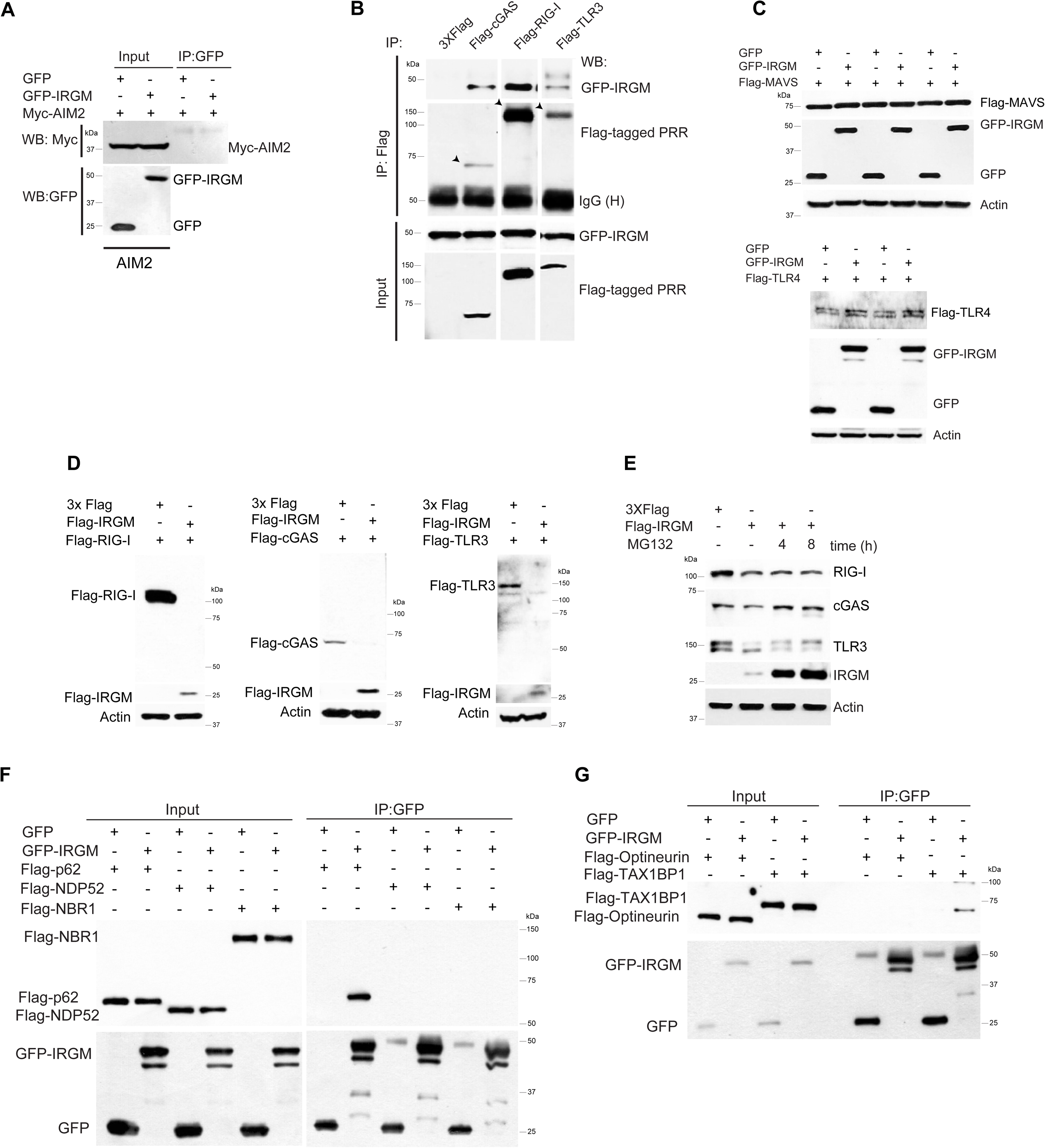
IRGM interacts and degrades cytoplasmic DNA and RNA sensor proteins to constrain IFN response. **(A)** Co-immunoprecipitation (Co-IP) analysis of interaction between GFP-IRGM and Myc-AIM2 in HEK293T cell lysates. **(B)** Co-IP analysis of the interaction between GFP-IRGM and Flag-cGAS, Flag-RIG-I and Flag-TLR3 in HEK293T cell lysates. **(C)** Western blot analysis with lysates of HEK293T cells transiently expressing GFP or GFP-IRGM and Flag-MAVS or Flag-TLR4. (n=3) **(D)** Western blot analysis of HEK293T cell lysates expressing 3X Flag or Flag-IRGM and Flag-RIG-I or Flag-cGAS or Flag-TLR3. **(E)** Control and HT29 cells stably expressing Flag IRGM were treated with MG132 (20 μM, 4 h and 8 h) and were subjected to immunoblotting with the indicated antibodies. (**F-G**) Co-IP analysis of the interaction between GFP IRGM with **(F)** Flag-p62 or Flag-NDP52 or Flag-NBR1 or (**G)** Flag-Optineurin or Flag-TAX1BP1 in HEK293T cell lysates.

**Supplementary Figure 5.**
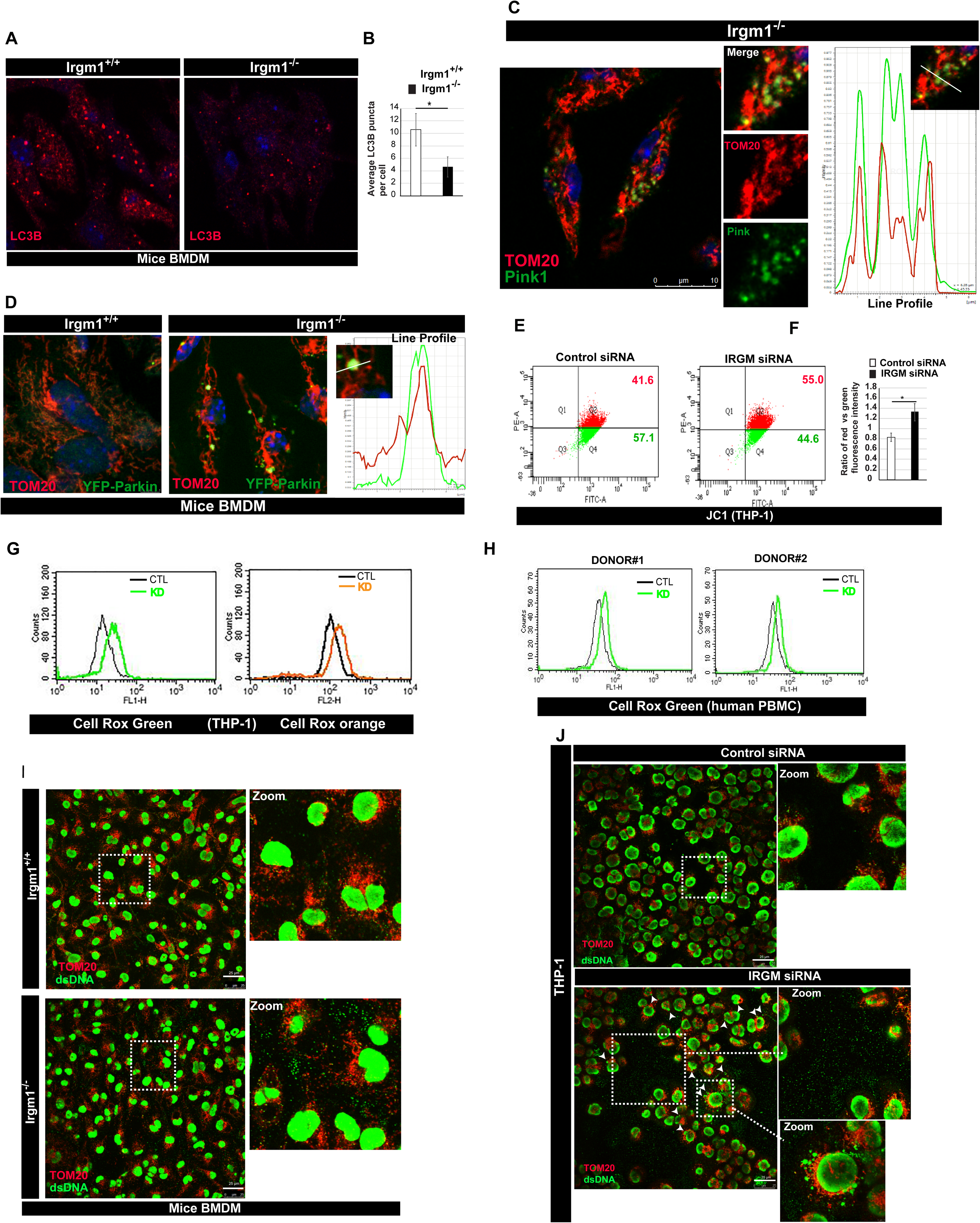
IRGM depletion results in defective mitophagy and enhanced cytosolic DAMPs. **(A)** Representative confocal images of *Irgm1^+/+^* and *Irgm1^-/-^* mice BMDMs immunostained with LC3B (red) antibody. The panels show LC3B puncta. **(B)** Bar diagram showing average LC3B puncta in *Irgm1^+/+^*and *Irgm1^-/-^* mice BMDMs. (n = 3, mean ± SD, ∗p ≤ 0.05, Student’s unpaired t-test). **(C)** Representative confocal images of *Irgm1^-/-^* mice BMDMs immunostained with TOM20 (red) and Pink1 (green) antibodies. Right Panel: Colocalization analysis using line intensity profiles. **(D)** Representative confocal images of *Irgm1^+/+^* and *Irgm1^-/-^* mice BMDMs transfected with YFP-Parkin (green) and immunostained with Tom20 (red) antibody. Right Panel: Colocalization analysis using line intensity profiles. **E)** Representative dot plot showing flow cytometry analysis of control and IRGM siRNA knockdown THP-1 cells stained with JC-1 dye (2 μM, 30 mins). **(F)** The graph depicts the ratio of red vs. green fluorescent intensity in control and IRGM siRNA knockdown THP-1 cells stained with JC-1 dye. Mean ± SD, n = 3, *P < 0.05, Student’s unpaired t-test. **(G)** Representative flow cytometry analysis of control and IRGM siRNA transfected THP-1 cells stained with CellRox green and CellRox orange dye (1μM, 30 min). (n=4). **(H)** Representative flow cytometry analysis of control and IRGM siRNA transfected human PBMC’s from two donor’s stained with CellRox green dye (1μM, 30 min). **(I)** Representative confocal images of *Irgm1^+/+^*and *Irgm1^-/-^* mice BMDMs immunostained with dsDNA (green) and TOM20 (red) antibodies. The *Irgm1^-^* zoom panels show extracellular DNA in cytoplasm. **(J)** Representative confocal images of control siRNA and IRGM siRNA transfected THP-1 cells immunostained with dsDNA (green) and TOM20 (red) antibodies. The zoom panels show extracellular DNA and cytoplasmic micronuclei. The white arrows in IRGM siRNA panel depict the micronuclei. The microscopic scales are as depicted in the images

**Supplementary Figure 6.**
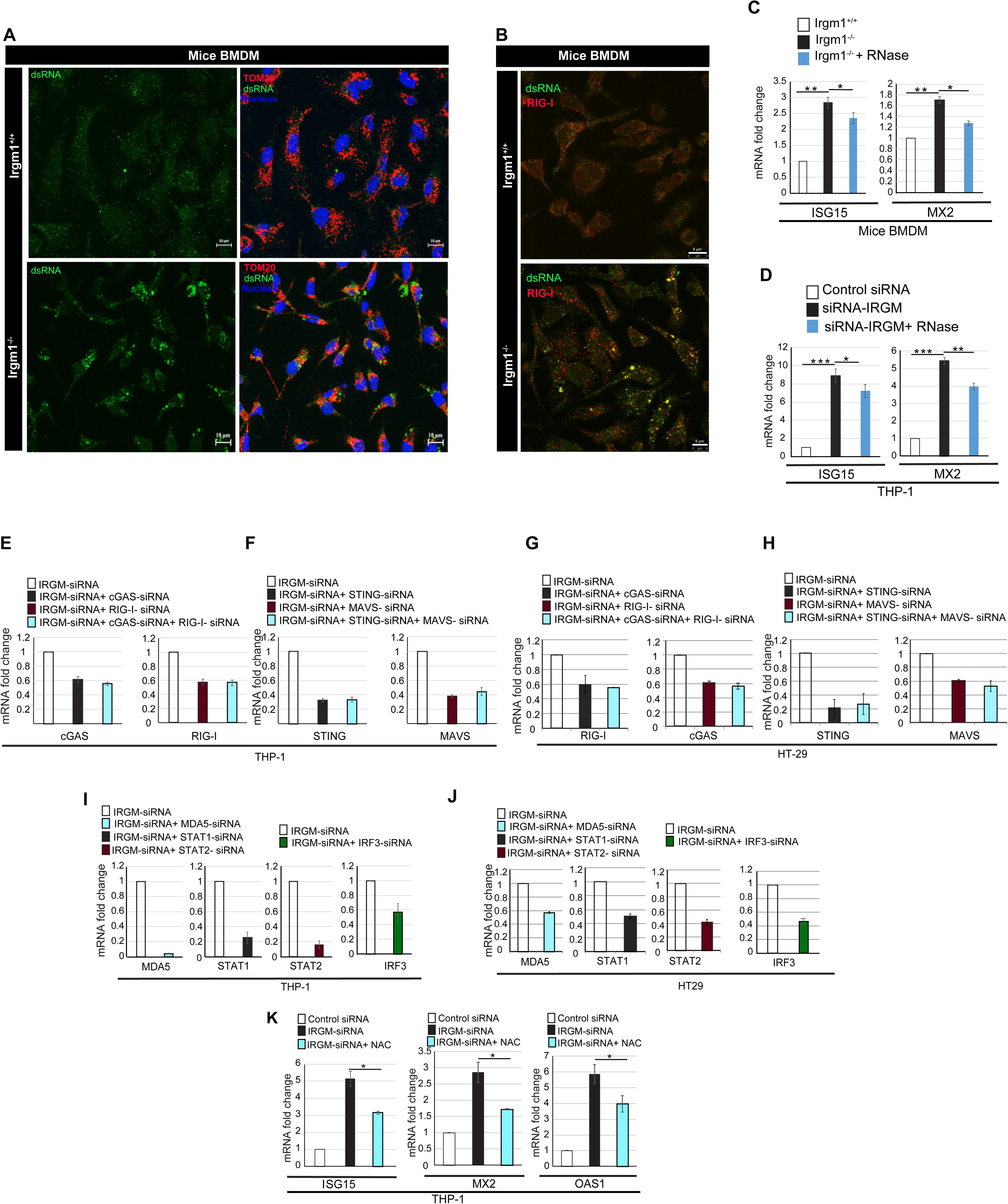
Cytosolic DAMPs in IRGM-depleted cells invoke nucleic acid-sensing pathway for activation of IFN response. **(A-B)** Representative confocal images of *Irgm1^+/+^* and *Irgm1^-/-^*mice BMDMs immunostained with dsRNA (green) and (**A**) TOM20 (red) or (**B**) Rig-I (red) antibodies. **(C)** The qRT-PCR analysis with RNA isolated from *Irgm1^+/+^* and *Irgm1^-/-^* mice BMDMs electroporated with RNase III and H (10 unit for 1hr) as indicated. (n = 3, mean ± SD, ∗p ≤ 0.05, ∗∗p ≤ 0.005, Student’s unpaired t-test). **(D)** The qRT-PCR analysis with RNA isolated from control and IRGM siRNA knockdown THP-1 cells electroporated with RNase III and H (10 unit for 1hr) as indicated. (n = 3, mean ± SD, ∗p ≤ 0.05, ∗∗p ≤ 0.005, ***p < 0.0005, Student’s unpaired t-test). **(E-F)** Control and IRGM knock down THP-1 cells were transfected with indicated siRNA and total RNA was subjected to qRT PCR with (**E**) cGAS and RIG-I primers (**F**) STING and MAVS primers. **(G-H)** Control and IRGM knock down HT-29 cells were transfected with indicated siRNA and total RNA was subjected to qRT PCR with (**G**) cGAS and RIG-I primers (**H**) STING and MAVS primers. **(I)** The qRT-PCR analysis with RNA isolated from control and IRGM siRNA knockdown THP-1 cells as indicated. **(J)** The qRT-PCR analysis with RNA isolated from control and IRGM siRNA knockdown HT29 cells as indicated. **(K)** qRT-PCR analysis with RNA isolated for control siRNA or IRGM siRNA transfected THP-1 cells untreated or treated with N-acetyl-L-cysteine (NAC, 1 mM, 2 h). Mean ± SD, n = 3, *P < 0.05, Student’s unpaired t-test.

**Supplementary Figure 7.**
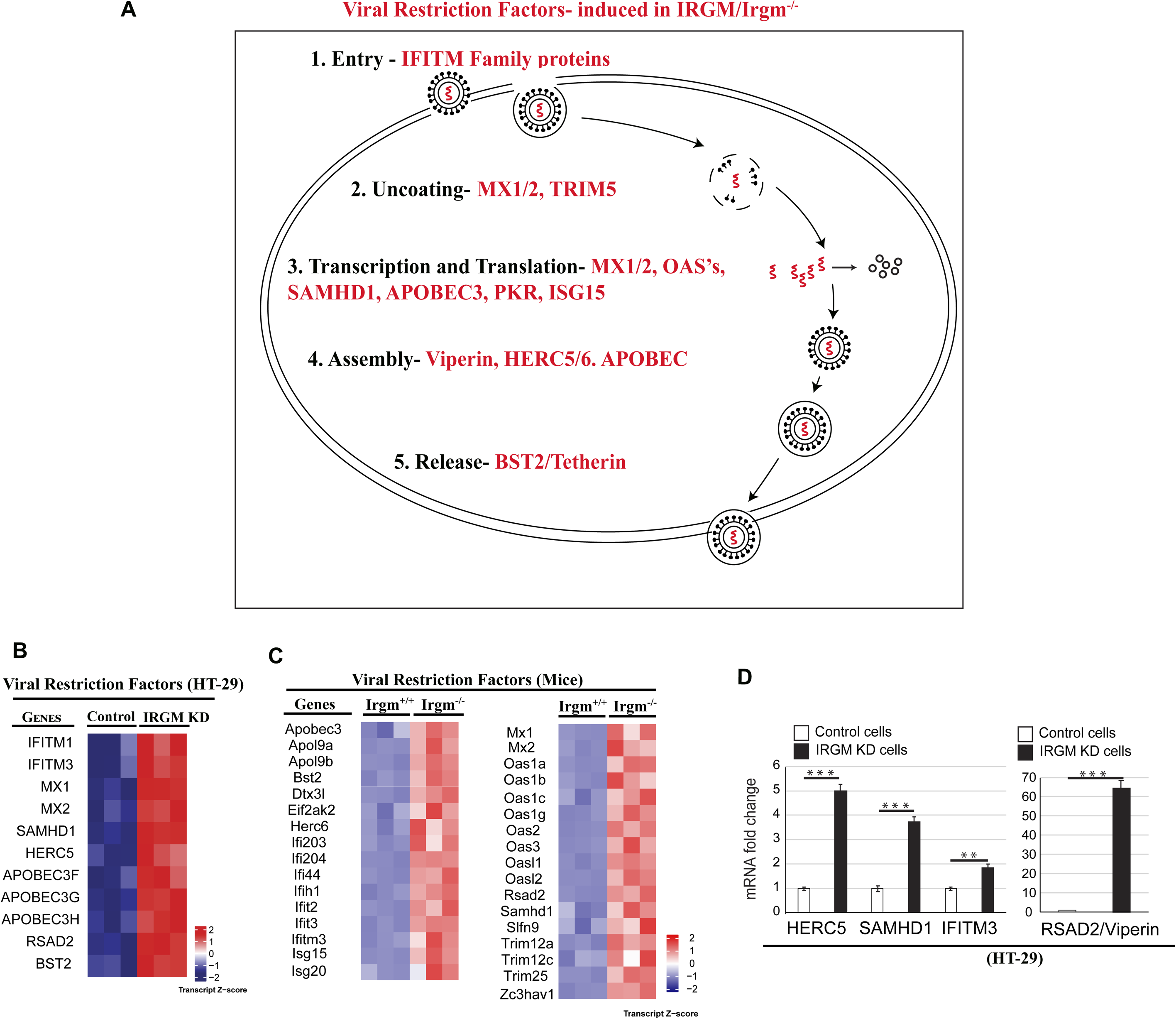
The depletion of IRGM invokes cell-intrinsic antiviral state. **(A)** Pictorial representation of stages (black font) of a typical life cycle of RNA viruses with host viral restriction factors (red font) induced in IRGM/Irgm1-depleted cells. **(B, C)** Heatmaps representing the expression pattern of viral restriction factors in (**A**) control and IRGM knockdown HT29 cells and (**B**) *Irgm1^+/+^* and *Irgm1^-/-^*mice. **(D)** The RNA isolated from control and IRGM knockdown HT29 cells and subjected to qRT-PCR with indicated viral restriction factor genes. Mean ± SD, n = 3, *p < 0.05, **p < 0.005, ***p < 0.0005, Student’s unpaired t-test.

## Materials and Methods

### Cell Culture

The cell lines, HT29 (ATCC #HTB-38), THP-1 (ATCC #TIB-202), HEK293T (ATCC #CRL-11268) were obtained from American Type Culture Collection (ATCC). Vero cells (African green monkey kidney epithelial cell line) was a kind gift from Dr. M.M. Parida, DRDE, Gwalior, India. HEK293T, HT29 and Vero cells were grown in DMEM medium (Gibco #10569044) supplemented with 10% fetal bovine serum (FBS) and penicillin/streptomycin (10,000 units/mL). Human monocytic cell line, THP-1 and BMDMs were grown in RPMI-1640 (Gibco#61870127) media supplemented with 10% FBS, 5 mM L-glutamine, glucose (5%), HEPES buffer, sodium pyruvate (1 mM), penicillin/streptomycin (10,000 units/mL). All the experiments were performed with cells before the 20^th^ passage was reached.

### Isolation of human peripheral blood mononuclear cells (PBMCs)

Isolation of human PBMCs was performed using density gradient centrifugation method using Histopaque(Sigma #10771) as per the manufacturer’s recommendation. Briefly, 5 mL of homogenized blood was mixed with an equal volume of DPBS and slowly layered on the 3 mL Histopaque. The sample was then centrifuged at 400 g for 20 minutes (min) at room temperature (RT) using slow acceleration and deceleration. Buffy coat generated in the middle of the gradient was collected, mixed with DPBS and centrifuged at 100 g for 10 min at RT. Cells were then washed twice with DPBS and re-suspended in RPMI-1640 supplemented with 10% FBS and incubated at 37°C with 5% CO_2_ overnight. Cells were counted in an Invitrogen countless II automated cell counter using an equal volume of trypan blue (Sigma #T8154). Total RNA was extracted using TRIZOL (Invitrogen #15596018).

### CRISPR Knockout Cell Lines

For generation of IRGM knockout in cell lines, HT-29 cells were transfected with an IRGM CRISPR/Cas9 KO plasmid (sc-407889) and IRGM HDR plasmid (sc-407889-HDR) using Lipofectamine 2000 DNA transfection reagents(Invitrogen #11668027). After 48 hour(h)of transfection, cells were treated with 2 μg/mL puromycin (InvivoGen #ant-pr-1) for 7 days. Single clones were selected, and IRGM knockout was confirmed by western blotting using anti-IRGM antibody.

### Lentivirus production and generation of stable IRGM knockdown cells

For generation of lentivirus, HEK293T cells cultured in 15 cm plate and transfected with control shRNA or IRGM shRNA (Santa Cruz #TRCN0000197363; Sequence: CCGGCCGGTATGACTTCATCATGGTCTCGAGACCATGATGAAGTCATACCGGTTTTTTG) plasmid (22.5 μg), the pMD2G plasmid (7.9 μg) and pCMVR 8.74 (14.6 μg) using CalPhos Mammalian Transfection Kit (Clonetech #631312). After 36 h of transfection, the viruses in culture medium were harvested and centrifuged at 500 g for 5 min at 4°C. Culture medium was then filtered through 0.22 µM filter and used for the transduction and generation of stable cell lines. For generation of stable IRGM knockdown cells, HT29 cells were plated in 6 well plates and transduced with virus particles. After 24 h, the medium was replaced and kept for another 48 h. IRGM stable knockdown cells were selected using 2µg puromycin for one week. IRGM knockdown was confirmed by western blotting using anti-IRGM antibody and qRT-PCR using IRGM TaqMan gene expression assay.

### Virus

Chikungunya virus (CHIKV) strain DRDE06 (accession no. EF210157.2) was gifted by Dr. M. M. Parida, DRDE Gwalior, India. Japanese encephalitis virus (JEV) strain GP78 (accession no. AF075723) was from Dr. A. Basu, NBRC, India and Herpes simplex virus-1(HSV-1) KOS strain (accession no.JQ673480.1) was kindly gifted by Dr. Roger Everett, Glasgow University.

### Virus infection

HT29 control and IRGM knockdown cells were seeded in 6 well plates. Next day, the cells were infected with either CHIKV or JEV or HSV-1 with a multiplicity of infection (MOI) 5 as described earlier (Kumar et al., 2014; Nayak et al., 2017). Briefly, the confluent monolayers were washed in sterile 1X PBS and infected with the virus (diluted in serum-free media) for 1.5 h with manual shaking at an interval of 10 min. Then, the cells were washed in sterile 1X PBS and were maintained in complete DMEM media in the incubator till harvest. The infected cells were collected for western blotting and qRT-PCR analysis.

### Plaque assay

The plaque assay was performed to quantitate the release of new infectious viral particles according to the previously mentioned protocol(Kumar et al., 2014). Briefly, virus-infected cell culture supernatants were serially diluted in serum-free media and confluent Vero cells seeded in 12 well cell culture plates were infected as per the protocol mentioned above. After infection, the cells were overlaid with methylcellulose (Sigma #M0387) containing DMEM and kept in the incubator for 3-4 days until visible plaques developed. Then the cells were fixed in 8% formaldehyde at RT, washed gently in tap water followed by staining with crystal violet. The number of plaques were counted manually in terms of plaque forming unit/mL (PFU/mL).

### Transient transfection with siRNA

The THP-1 cells were electroporated (Neon, Invitrogen #MPK5000; setting: 1400V, 10ms, 3 pulses using 100 µL tip#MPK10096) with non-targeting siRNA (30 nM) or specific siRNA (30 nM) and incubated for 24h. Another round of siRNA transfection was performed after 24 h in similar condition as described above and incubated for next 48 h. The siRNA (10 nM) transfection in HT29 cells were performed using the Lipofectamine® RNAiMAX (Invitrogen #13778075)as per the manufacturer’s instructions. Following siRNA were used in the present study: Non-specific siRNA (SMARTpool: siGENOME ns siRNA; Dharmacon #D-001206-13-20),IRGM siRNA (SASI_HS02_00518571), p62 siRNA (SASI_Hs01_00118616), BECLIN1 siRNA (SASI_Hs02_00336256), Human RIG I siRNA (SASI_Hs01_00047980), TLR3 siRNA (SASI_Hs01_00231802), Human cGAS siRNA (SASI_Hs01_00197466), MAVS siRNA (SASI_Hs01_00128708), MDA5 siRNA (SASI_Hs01_00171929), STING siRNA (SASI_Hs02_00371843), TRIF siRNA (SASI_Hs01_00226929), IRF3 siRNA (SASI_Hs02_00332144), STAT1 siRNA (SASI_Hs02_00343387), STAT2 siRNA (SASI_Hs01_00111824), Mice RIG I siRNA (SASI_Mm01_00145086), Mice cGAS siRNA (SASI_Mm01_00129826)

### Transient transfection with plasmids

For transient expression, HEK293T cells were transfected with plasmids using the calcium phosphate method. THP-1 cells were transfected using Neon electroporation system with the following parameters: 1300V, 30ms, 1pulse. Briefly, 2x10^6^ THP-1 cells were transfected with p62(30nM) or BECLIN1 (30nM) siRNA. After 72h, the cells were transfected with EGFP (3μg) or GFP-IRGM (3 μg) or 3X flag or flag-IRGM (3 μg). After 4h, cell lysates were prepared and western blotting was performed.

### DNase I Transfection in THP-1 cells

Approximately 3x10^6^ THP-1 cells were electroporated (Neon, Invitrogen; setting: 1400V, 10ms, 3 pulses using the 100 µL tip) with non-targeting siRNA (30nM) or IRGM siRNA (30nM). After 24h the cells were again transfected with siRNAs. Next day, the cells were electroporated (1400V, 10ms, 2 pulses using the 100 µL tip) with either 10 µg or 5ug BSA (as control) or 10 µg or 5 µg DNAse I enzyme (NEB #M0303S). The cells were harvested after indicated time points and analyzed by qRT-PCR.

### RNase III and RNase H Transfection in THP-1 cells

Approximately 3x10^6^ THP-1 cells were electroporated (Neon, Invitrogen; setting: 1400V, 10ms, 3 pulses using the 100 µL tip) with non-targeting siRNA (30nM) or IRGM siRNA (30nM). After 24h the cells were again transfected with siRNAs. Next day, the cells were electroporated (1400V, 10ms, 2 pulses using the 100 µL tip) with either 10 µg BSA (as control) or 10 unit of RNase III (NEB #M0245S) and RNase H (NEB #M0297S) enzymes each. The cells were harvested after 1h and analyzed by qRT-PCR.

### Sample preparation for RNA-Sequencing

RNA was extracted from brain tissues and BMDMs of three different mice and HT29 cells(three biological replicates) using RNeasy mini kit (QIAGEN, #74104). The quality and quantity of total RNA was checked using agarose gel and Qubit 3.0. After assessing the quality of RNA, 800-900 ng of total RNA was subjected to NEBNext® ultra™ directional RNA library prep kit for Illumina® (#E7420) using NEBNextPoly(A) mRNA magnetic isolation module (NEB #E7490). The QC of the prepared library was performed using High-Sensitivity Tape Station Kit (Agilent 2200, 5067-5585 (reagents) #5067-5584 (screen tapes)) for fragment length distribution and Qubit dsDNA HS assay kit (Invitrogen, Q32851) for quantifying the library. The library was sequenced using the HiSeq 4000 Illumina platform.

### RNA-sequencing data processing and gene expression analysis

Paired-end (PE) reads quality checks were performed using the FastQC v.0.11.5 (http://www.bioinformatics.babraham.ac.uk/projects/fastqc/). The adaptor sequence was trimmed using the ‘bbduk’ using BBDuk version 37.58, using ‘bbduk.sh’ script. The files were further processed for alignment using STAR v.2.5.3a with default parameters (Dobin et al., 2013). The mouse genome build GRCm38.p6genome.fa, was used from Gencode.vM18.annotation.gtf. Duplicates were removed from Picard-2.9.4 (https://broadinstitute.github.io/picard/) from the aligned bam files. Count matrix for each comparison was generated for differential gene analysis, featureCounts v.1.5.3 from subread-1.5.3 package (http://bioinf.wehi.edu.au/featureCounts/) was used with Q = 10 for mapping quality, and these count files were used as input for downstream differential gene expression analysis with DESeq2 version 1.14.1 (Love et al., 2014). Genes with read counts of ≤ 10 in any comparison were removed followed by using DESeq2 ‘R’ library for count transformation and statistical analysis. ‘P’ values were adjusted using the Benjamini and Hochberg multiple testing correction(Haynes, 2013). Significantly differentially expressed genes were identified based on a fold-change of 1.5-fold or greater (up- or down-regulated) and a p-value less than 0.05.

### Clustering and heatmap generation

Unsupervised hierarchical clustering was performed and the heatmap was plotted using ‘ComplexHeatmap’ library using ‘R’ Bioconductor package (https://www.bioconductor.org/) where the gene expression matrix was transformed into z-score(Gu et al., 2016).

### Western Blotting

Cell and tissue lysate preparation and western blot analysis was performed as described previously (Mehto et al., 2019). Briefly, cell lysates were prepared using the NP-40 lysis buffer (Invitrogen #FNN0021) containing protease inhibitor cocktail (Roche #11836170001), phosstop (Roche #4906845001) and 1mM PMSF (Sigma #P7626). Mice tissues (100 mg) were homogenized in 1 mL Radio-immunoprecipitation assay (RIPA) buffer (20 mM Tris, pH 8.0; 1mM, EDTA; 0.5 mM, EGTA; 0.1%Sodium deoxycholate; 150 mM NaCl; 1% IGEPAL; 10% glycerol) with protease inhibitor cocktail, phosstop and 1mM PMSF using tissue tearor (BioSpec #985370). Lysates were separated using SDS-polyacrylamide gel, transferred onto nitrocellulose membrane (Bio-Rad) and blocked for 1 h in 5% skimmed milk followed by incubation in primary antibody overnight at 4°C. Membranes were then washed thrice with1X PBS/PBST and incubated for 1 h with HRP conjugated secondary antibody. After washing with PBS/PBST the blots were developed using enhanced chemiluminescence reagent (Thermo Fisher #32132X3).

### Mitochondrial isolation

Mitochondria from control and IRGM knockdown THP-1 cells were isolated using Qproteome mitochondria isolation kit (QIAGEN #37612) according to manufacturer’s instructions. Briefly, 10 x 10^6^ cells were lysed in ice cold lysis buffer (supplemented with protease inhibitor) by centrifuging at 1000g at 4°C for 10 min. The cell pellet was re-suspended in disruption buffer supplemented with a protease inhibitor. The cells were completely disrupted and the lysates were centrifuged at 1000 g for 10 min at 4°C. The supernatant was transferred and centrifuged at 6000 g for 10 min at 4°C to form a pellet containing mitochondria. The pellet was washed and re-suspended in a mitochondrial storage buffer for further analysis.

### Semi-denaturing Agarose Gel Electrophoresis

For MAVS oligomerization assay, the crude mitochondrial pellet was resuspended in 4X sample buffer (0.5X TAE, 10% glycerol, 2% SDS and 0.0025% bromophenol blue) followed by incubation at RT for 5 min and then loaded onto a vertical 1.5% agarose gel. Electrophoresis was performed in the running buffer (1X TAE and 0.1% SDS) for 40 min with a constant voltage of 100V at 4°C. After electrophoresis, the proteins were transferred onto a PVDF membrane for immunoblotting.

### Antibodies and dilution

Primary antibodies used in western blotting with dilutions: Flag (Sigma #F1804; 1:1000), c-Myc (Santa Cruz sc40; 1:750 and sc764; 1:1000), p62 (BD #610832; 1:2000), Actin (Abcam #ab6276; 1:5000), HA (CST #3724; 1:1000), GFP (Abcam #ab290; 1:5000), IRGM antibody rodent specific (CST #14979; 1:1000), IRGM (Abcam #ab69494; 1:500), RIG I (CST #3743; 1:1000), TLR3 (CST #6961; 1:1000), Cgas (CST #15102; 1:1000, Santa Cruz #sc515777; 1:500), STING (CST #3337, #13647; 1:1000), MDA5 (Santa Cruz #sc48031; 1:500), TRIF (Abcam #ab13810; 1:5000), MAVS (CST #3993; 1:1000, Santa Cruz #sc365334; 1:500), TBK1/NAK (CST #3504; 1:1000), pTBK1 (CST #5483S; 1:1000), pIRF7 (CST #12390S; 1:1000), IRF7 (CST #4920; 1:1000, Abcam #ab62505; 1:5000), pIRF3 (CST #29047; 1:1000), IRF3 (CST #11904; 1:1000, Abcam #ab25950; 1:5000) pSTAT1 (CST #7649; 1:1000), STAT1 (CST #9172; 1:1000), pSTAT2 (CST #88410; 1:1000, Abcam #ab53132; 1:5000) STAT2 (CST #72604; 1:1000), TLR4 (Santa Cruz #sc293072; 1:500), IRF9 (Abcam #ab51639; 1:5000), TLR7 (Santa Cruz #sc57463; 1:500), BECLIN1 (CST #3738; 1:1000), MX1 (CST #37849; 1:1000), OAS1 (CST #14498; 1:1000), ISG15 (CST #2743; 1:1000), SAMHD1 (CST #12361; 1:1000), BST2 (CST #19277; 1:1000), Viperin (CST #13996; 1:1000), APOBEC3G (CST #43584; 1:1000), AIM2 (CST #12948; 1:1000), NLRC4 (CST #12421; 1:1000), ph2AX (CST #2577; 1:1000), p21 (CST #2947; 1:1000), CARDIFF (Adipogen #SG-20B-0004-C100; 1:1000), C3d (R&D Systems #AF2655;1:2000), GAPDH (CST #2118; 1:1000), Normal rabbit IgG (CST #2729), LC3B (Sigma #L7543; 1:1000), Caspase1+p10+p12 (Abcam #ab179515; 1:1000), TOM20 (Santa Cruz #sc11415; 1:1000), PINK1 (CST #6946; 1:1000), PINK1 (Abcam #ab186303; 1:1000) Parkin (CST #4211; 1:1000), PARP1 (CST #9542; 1:1000), pMLKL (CST #37333; 1:1000), MLKL (CST #37705; 1:1000)

HRP conjugated secondary antibodies were purchased from Santa Cruz (1:2000) or Promega (1:5000) or Abcam (1:10000) or Novus (1:5000).

Primary antibodies used in immunofluorescence assays with dilutions: IRGM (Abcam # 69494; 1:100), dsRNA (Kerafast # ES2001; 1:60), TOM20 (Santa Cruz #sc11415; 1:150), RIG I (CST #3743, 1:50), MDA5 (Santa Cruz #sc48031; 1:50), TIA1 (Abcam #140595; 1:100), dsDNA (Abcam #ab27156; 1:10,000), Lamin A/C (Santa cruz #sc6215; 1:50), MX1 (CST #37849; 1:200), cGAS (CST #15102; 1:50 or Santa Cruz # sc515777; 1:50), PINK1 (Abcam #ab186303; 1:100), Parkin (CST #4211; 1:100)

### Co-Immunoprecipitation

Co-Immunoprecipitation assays were performed as described previously (Jena et al., 2018). The cells were lysed in NP-40 lysis buffer (Thermo Fisher #FNN0021) supplemented with protease inhibitor/phosphatase inhibitor cocktails and 1M PMSF for 30 min at 4°C and subjected to centrifugation. The supernatant was incubated with the desired antibody at 4°C (2 h to overnight) on rotational cell mixer followed by incubation with protein G dynabeads (Invitrogen, #10004D) for 2 h at 4°C. The beads were washed with lysis buffer and ice cold PBS(3-4 times) and the proteins were eluted by boiling for 5 min in 2X laemmlibuffer and proceeded for western blot analysis.

### Immunofluorescence Analysis

Approximately 10^5^ cells were seeded on coverslip and allowed to adhere to the surface. For THP-1, cells were differentiated into the macrophage-like state by addition of 50 ng/ml of phorbol 12-myristate 13-acetate (PMA) (Sigma #P8139) for 16 h. Next, the culture medium was replaced and incubated for 48 h. The adhered cells were fixed in 4% paraformaldehyde for 10 min and permeabilized with 0.1% Triton X-100 for 10 min, followed by blocking with 1% BSA for 30 min at RT. The cells were then incubated with primary antibody for 1 h at RT, washed thrice with 1X PBS, followed by 1 h incubation with AlexaFluor-conjugated secondary antibody. Cells were again washed thrice with 1X PBS, mounted (Prolong gold antifade, Invitrogen #P36931) and visualized using Leica TCS SP5 and TCS SP8STED confocal microscope. For Mitotracker CMXRos (Invitrogen #M7512) staining, cells were stained with the dye (10 nM, 30 mins) as per the manufacturer’s instructions, washed, fixed with 4% PFA for 10 min and visualized using Leica TCS SP5 confocal microscope.

### Luciferase assay

Luciferase assay was performed using a dual luciferase assay kit as per the manufacturer’s protocol (Promega #E1910). Briefly, HEK293T cells were cultured in 24 well plates and transfected with pISRE-luciferase reporter plasmid (100 ng) and renilla luciferase plasmid (50 ng) along with required plasmids (300 ng). After 36 h, growth medium was completely removed and after washing thrice with 1X PBS, cells were lysed using 100 µl 1X passive lysis buffer by incubating the plate on orbital shaker for 15 min at RT. Cell lysates were cleared by centrifugation for 30sec at 12000 g in a refrigerated centrifuge. In 96 well plate, 20 µL cell lysate mixed with 100 µL LARII reagents and luminescence was measured using PerkinElmer VICTOR Nivo multimode plate reader. Stop &Glo reagent was used to measure renilla luciferase activity.

### RNA Isolation and Quantitative Real-time PCR

RNA was extracted using TRIZOL by following the manufacturer’s protocol. The cDNA was synthesized using the high capacity DNA reverse transcription kit (Applied Biosystems,#4368813) and qRT-PCR was performed using Taqman master mix (Applied Biosystems, #4369016) or Power SYBR green PCR master mix (Applied Biosystems, #4367659) according to manufacturer’s protocol. For normalization of the assay, the housekeeping gene GAPDH or β-Actin was used. The fold-change in expression was calculated by the 2^−ΔΔCt^ method. Primer sequence for qRT PCR are as follows:

qRT primers:

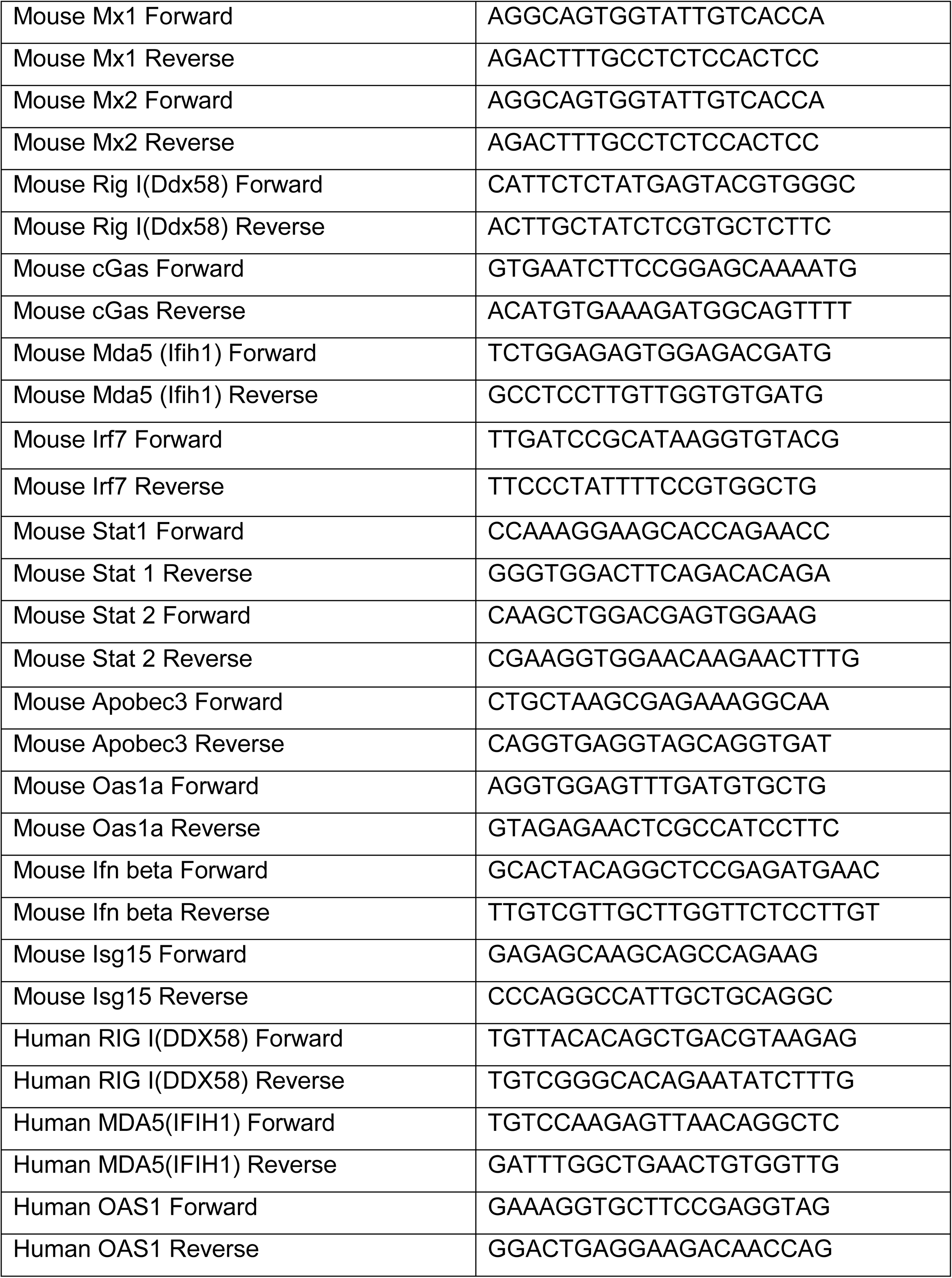
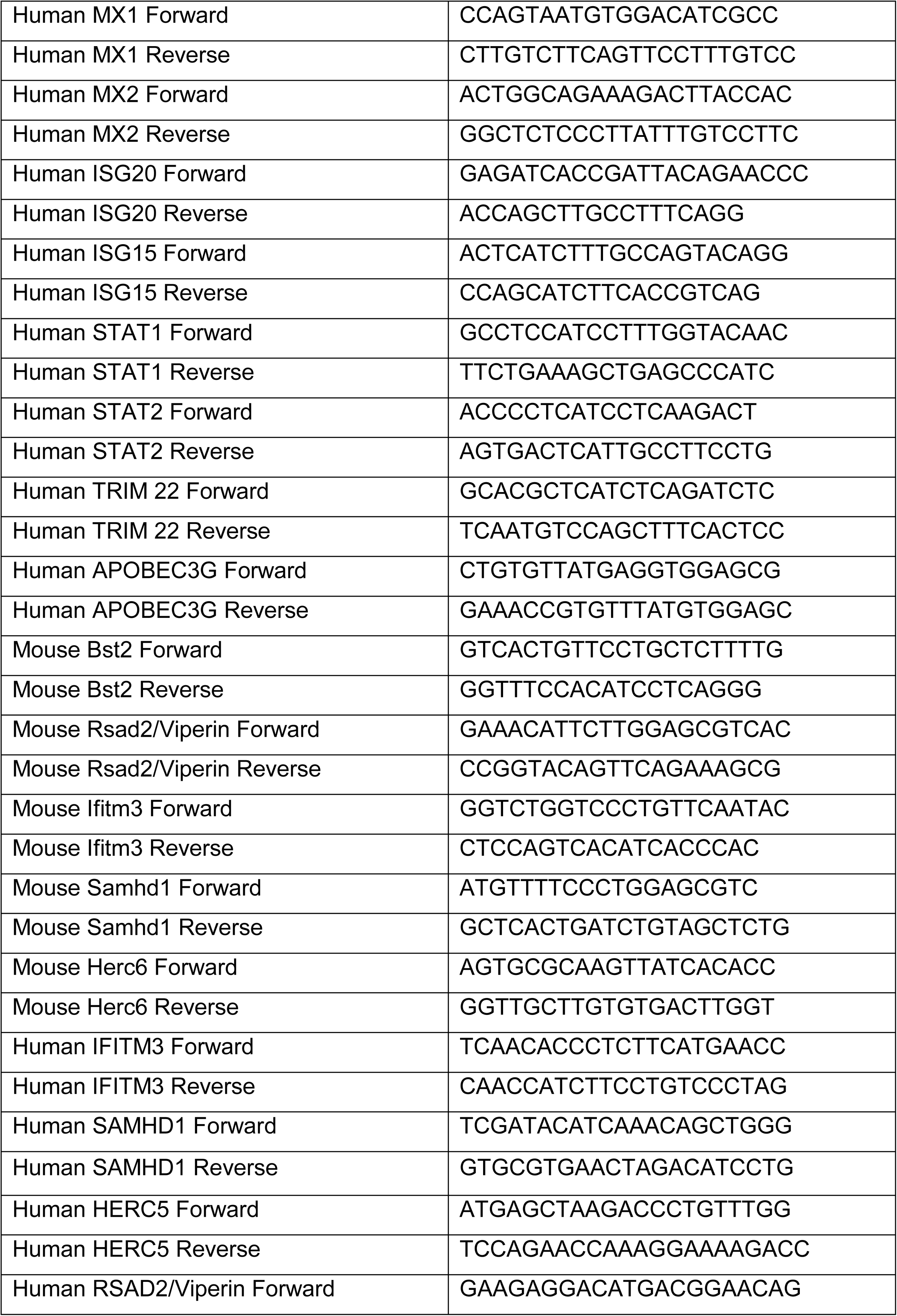
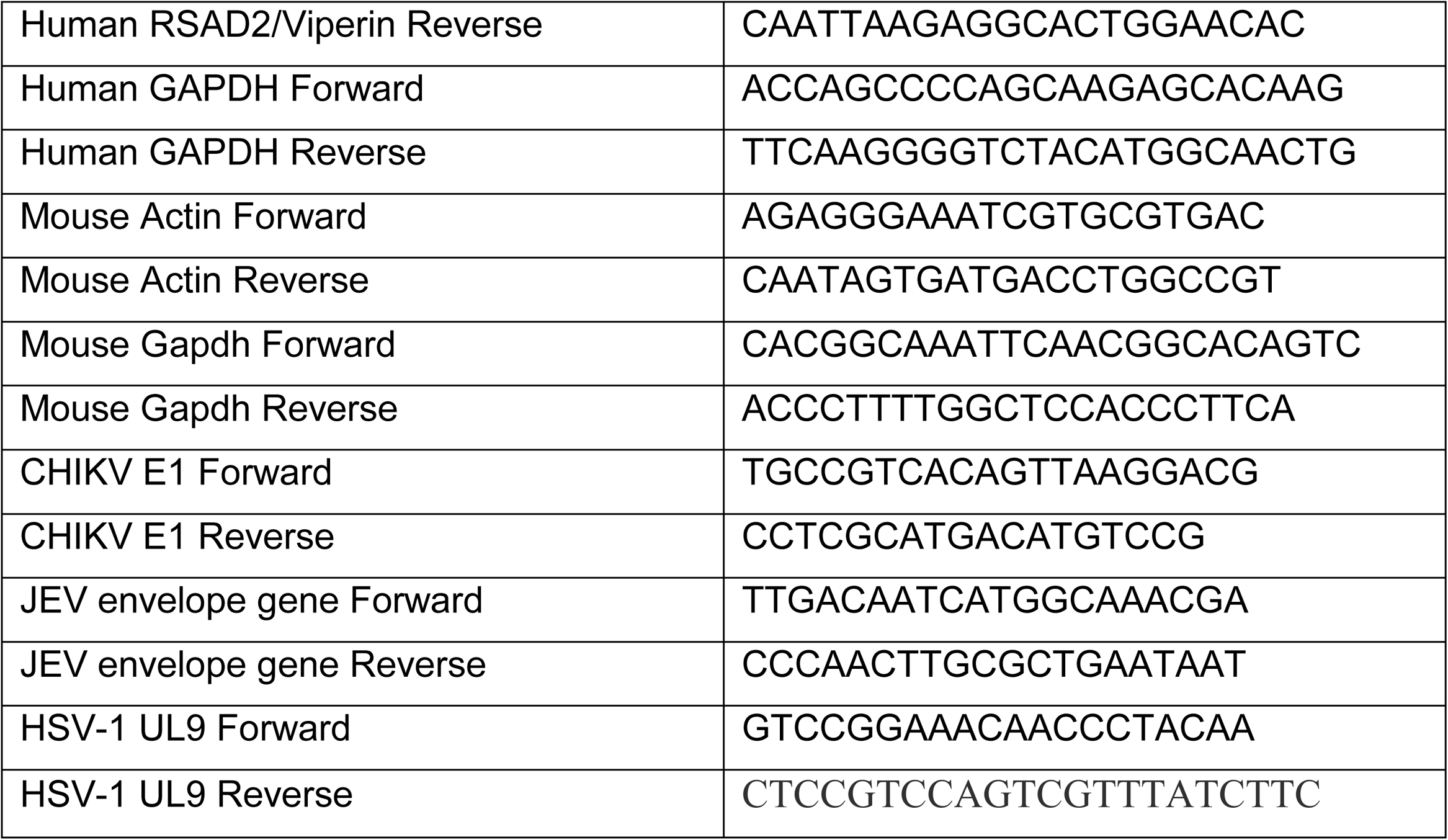

### Flow Cytometry

Mitochondrial membrane potential/polarization experiment was performed in live cells. The cells were incubated with CMXRos dye (Invitrogen #M7512) (10 nM, 30 min) or JC-1 (eBiosciences #65085138) (2 μM, 30 min) to measure the change in mitochondrial potential. Mitochondrial superoxide generation was measured using MitoSox (Invitrogen #M36008) staining (1μM, 20 min). The cellular ROS levels were measured using CellRox (Invitrogen #C10448) staining (1μM, 30 min). Briefly, 10^5^ cells were stained with the mentioned dyes and incubated for indicated time points at 37°C and 5% CO_2_. Samples were washed twice with 1XPBS and resuspended in 1X PBS, followed by acquisition on FACS Calibur (Beckton and Dickinson). The results were analyzed using the Cell Quest Pro software.

### Mice experiments

*Irgm1* knockout (C57BL/6) mice (*Irgm1^-/-^*) were maintained as described previously (Liu et al., 2013). The mice experiments were performed with procedures approved by institutional animal ethical committee (IAEC) at Institute of Life Sciences, Bhubaneswar, India. For each experiment, littermates were used, and their gender and age of mice were matched. The six days old *Irgm1* wild type (*Irgm1^+/+^*) and knockout mice pups (n=12, three experiments) were infected with DRDE-06 strain of Chikungunia virus (1x10^7^ PFU/mouse) through intradermal route. Mice were monitored daily for muscle weakness, flaccid paralysis symptoms and changes in the body weights. The paralysis scores were assigned as follows: normal; 0, single leg paralysis; 1, both leg paralysis; 2, severe paralysis; 3. The mice were sacrificed and different organs were collected and processed further for qRT-PCR analysis.

### Mice Bone marrow cells isolation and differentiation into macrophages

The bone marrow cells from wild type (*Irgm1^+/+^*) and knockout (*Irgm1^-/-^*) mice were isolated and differentiated into macrophages by standard procedure. Briefly, six to eight weeks old male C57BL/6 *Irgm1^+/+^*and *Irgm1^-/-^* mice were sacrificed by cervical dislocation, bone marrow cells from the tibia and femurs were flushed out in RPMI medium. Red blood cells were removed by cell lysis buffer containing (155 mM NH_4_Cl, 12 mM NaHCO_3_ and 0.1 mM EDTA). Bone marrow cells were differentiated in RPMI medium (10% FBS, 1mM sodium pyruvate and 0.05 M 2-mercaptoethanol) containing 20 ng/mL mouse M-CSF (Gibco #PMC2044) for 5 days. On every alternate day, media was replaced with fresh media containing M-CSF.

